# A low-cost, thermostable, cell-free protein synthesis platform for on demand production of conjugate vaccines

**DOI:** 10.1101/2022.08.10.503507

**Authors:** Katherine F. Warfel, Asher Williams, Derek A. Wong, Sarah E. Sobol, Primit Desai, Jie Li, Yung-Fu Chang, Matthew P. DeLisa, Ashty S. Karim, Michael C. Jewett

**Author notes:** To whom correspondence should be addressed, Michael C. Jewett, **Email:**.

## Abstract

Cell-free protein synthesis systems that can be lyophilized for long-term, non-refrigerated storage and transportation have the potential to enable decentralized biomanufacturing. However, increased thermostability and decreased reaction cost are necessary for further technology adoption. Here, we identify maltodextrin as an additive to cell-free reactions that can act as both a lyoprotectant to increase thermostability, as well as a low-cost energy substrate. As a model, we apply optimized formulations to produce conjugate vaccines for ~$0.50 per dose after storage at room temperature or 37 °C for up to 4 weeks and ~$1.00 per dose after storage at 50 °C for up to 4 weeks. We show that these conjugates generate bactericidal antibodies against enterotoxigenic *E. coli* (ETEC) O78 O-polysaccharide, a pathogen responsible for diarrheal disease, in immunized mice. We anticipate that our low-cost, thermostable cell-free glycoprotein synthesis system will enable new models of medicine biosynthesis and distribution that bypass cold-chain requirements.

## Introduction

Synthetic biology promises to transform planet and societal health by producing energy, materials, fuels, foods, medicines, and more^1–3^. Unfortunately, current state-of-the-art biomanufacturing practices require expensive, centralized facilities to grow cells used to make bioproducts, tend to be inflexible because of the cost of customization, and can require cold-chain for distribution (e.g., mRNA vaccines)^4,5^.

Cell-free gene expression (CFE) systems have recently matured as an approach to address these limitations^6–14^. The foundational principle is that biological processes (e.g., protein biosynthesis, metabolism) can be conducted outside of living cells in crude cell-free lysates^15,16^. Key features of CFE systems include that they are (i) distributable through freeze drying^17^, which allows simple distribution before rehydration at the point-of-use^12,13,18–24^, (ii) scalable linearly from 1-nL to 100-L^25,26^, which accelerates process development, and (iii) do not require unique production cell lines for each product, which facilitates rapid customization and product switching^10,12^. Taken together, these features have the potential to advance new paradigms in decentralized manufacturing. For example, lyophilized cell-free systems have already been used to manufacture a variety of products in a manner suitable for portable biomanufacturing (e.g., conjugate vaccines^13^, erythropoietin^10^, and granulocyte-macrophage colony-stimulating factor^11^).

While recent breakthroughs in freeze-dried CFE systems have set the stage for creating a disruptive, distributed protein biosynthesis technology, adoption of CFE systems remains limited by costs and thermostability. For example, we recently developed a modular, *in vitro* conjugate vaccine expression (iVAX) platform that can be freeze-dried and rehydrated for decentralized production of conjugate vaccines^13^. However, lyophilized iVAX reactions are on the order of ~$5.00 per reaction and are not stable at elevated temperatures, making them infeasible for distribution and use in resource-limited settings. The Meningitis Vaccine Project recently benchmarked parameters for conjugate vaccine distribution in remote settings with the WHO approval of the MenAfriVac vaccine for controlled temperature chain storage for 4 days at up to 40 °C at <$0.50 per dose^27,28^. Adjusting CFE reaction formulations could address these challenges in our cell-free conjugate vaccine production platform. However, to date, CFE optimizations have typically sought to address either cost^29–32^ or thermostability^33–35^ rather than considering both formulation properties together.

In this work, we set out to address both cost and stability of CFE reactions together, to identify a low-cost and thermostable formulation for decentralized manufacturing. As a model, we selected production of conjugate vaccines, which are among the most effective methods for preventing bacterial infections that are predicted to threaten up to 10 million lives by 2050^36–40^. First, we screened sugar additives that could potentially serve as both lyoprotectant and energy system. We identified maltodextrin as the best lyoprotectant. We then optimized the formulation to also use maltodextrin as a low-cost energy substrate, reducing the reaction cost ~4-fold and providing thermostability of lyophilized reactions after 4 weeks of storage at room temperature, 37 °C, and 50 °C. Finally, we demonstrated that cell-free glycoprotein synthesis machinery is still active in all formulations under these storage conditions by producing relevant and effective anti-diarrheal conjugate vaccine molecules (ETEC O78 O-antigen conjugated to the approved carrier PD) for as low as ~$0.50 per dose.

## Results and Discussion

### Maltodextrin enhances thermostability of CFE reactions

With the goal of decreasing costs and increasing stability of CFE reactions, we first benchmarked the thermostability of our CFE formulation using a common protein expression lysate derived from BL21 Star (DE3) cells. We lyophilized 5-μL CFE reactions containing all reagents for the PANOx-SP-based system^41^, which uses the phosphorylated secondary energy substrate phosphoenolpyruvate (PEP), supplemented with DNA encoding superfolder green fluorescent protein (sfGFP). Then, after one, two, and four weeks of storage at 37 °C in vacuum sealed bags with desiccant cards, we rehydrated lyophilized reactions with 5 μL of water and measured sfGFP concentrations via fluorescence (**Figure 1A**). Rehydrated controls (zero-week timepoint) produced protein comparable to controls that were never lyophilized (fresh) (**Figure S1**), but lyophilized CFE reactions with no lyoprotectant additives did not produce sfGFP after 1 week of storage at 37 °C (**Figure 1B**; black circles). Consistent with previous studies^35^, these data indicated that lyophilized one pot CFE reactions are not thermostable at elevated temperatures.

**Figure 1.**
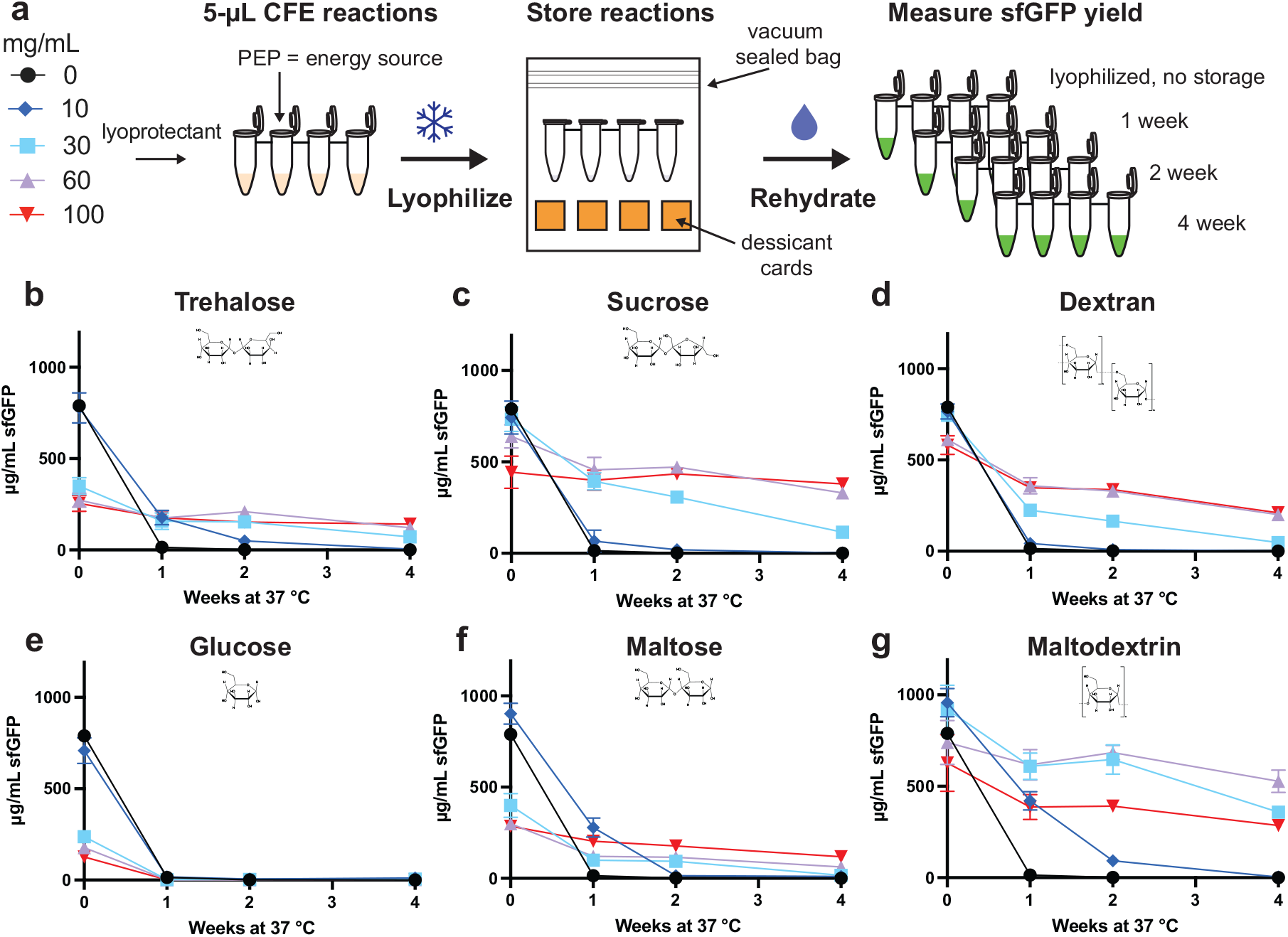
Maltodextrin enhances stability of cell-free gene expression (CFE) reactions stored at 37 °C. (A) Schematic of CFE reaction set-up and lyophilization for the screening of lyoprotectants. The impact on sfGFP production of (B) trehalose, (C) sucrose, (D) dextran, (E) glucose, (F) maltose, and (G) maltodextrin at concentrations of 0 mg/mL in black circles, 10 mg/mL in blue diamonds, 30 mg/mL in light blue squares, 60 mg/mL in purple triangles, and 100 mg/mL in inverted red triangles on lyophilized CFE reactions after storage was measured. Reactions were rehydrated with 5 μL of water and incubated at 30 °C for 20 hours after one, two, and four weeks of storage at 37 °C. Error bars represent standard deviation of 3 CFE reactions.

We next sought to identify low-cost lyoprotectant additives that could confer storage stability at elevated temperatures (37 °C). Specifically, we explored the use of trehalose^34,42^, sucrose^43^, and dextran^35^, which have previously been shown to enhance lyophilized reaction stability (**Figure 1B-D**). In addition, we wanted to test whether sugars that have been demonstrated as low-cost, secondary energy sources in CFE systems, such as glucose^31,44,45^, maltose^46,47^, maltodextrin^46–49^, could also protect or stabilize reactions during lyophilization and storage (**Figure 1 E-G**).

We supplemented CFE reactions with 0 to 100 mg/mL of each lyoprotectant individually prior to lyophilization. No significant loss in activity was observed from the lyophilization process, although some lyoprotectants (e.g., trehalose) were detrimental to protein yields (**Figure S1**). Then, after one, two, and four weeks of storage, we rehydrated lyophilized reactions with 5 μL of water and measured sfGFP concentrations via fluorescence. The addition of trehalose, glucose, and maltose each significantly decreased protein expression at concentrations greater than 10 mg/mL in fresh and lyophilized reactions (**Figure S1; Figure 1B, E,** and **F**). Supplementing reactions with 100 mg/mL sucrose or dextran retained ~85% and ~36% of the freshly lyophilized reaction activity (zero-week timepoint for each condition) after four weeks of storage, respectively (**Figure 1C** and **D**). However, adding just 60 mg/mL of maltodextrin retained 71 ± 18% of the freshly lyophilized reaction activity (zero-week timepoint for this condition) and achieved higher overall protein yields after 4 weeks of storage (528 ± 61 μg/mL sfGFP) than sucrose-protected reactions (**Figure 1G**). Of note, adding maltodextrin protects CFE reactions without any additional costly additives such as DMSO or stabilizers^35^ resulting in a simplified and more cost-effective solution.

### Maltodextrin can be used as a low-cost CFE lyoprotectant and energy source

After identifying that maltodextrin could be used as an effective lyoprotectant, we wanted to explore whether this polysaccharide could simultaneously preserve the reaction and act as an energy source for CFE reactions. Maltodextrin, a non-phosphorylated substrate, with the addition of exogenous phosphate, can be broken down into early glycolytic intermediates and used to fuel protein synthesis^47–49^. By having a dual-use for maltodextrin (~$0.02 per mL reaction with 60 mg/mL maltodextrin) and replacing PEP in the PANOx-SP system, we could potentially reduce the cost of CFE reagent formulation from ~$4.93 per mL reaction to ~$2.89 per mL of reaction (a 59% reduction) (**Table S1-3; Figure 2A**). Further, replacing nucleotide triphosphates (NTPs) with nucleotide monophosphates (NMPs), which can be phosphorylated in the cell-free reaction, and removing non-essential additives like tRNA and CoA^29–31^ could yield a minimal formulation (MD min) costing ~$1.38 per mL of CFE reaction, a quarter of the cost per mL of the PANOx-SP CFE system.

**Figure 2.**
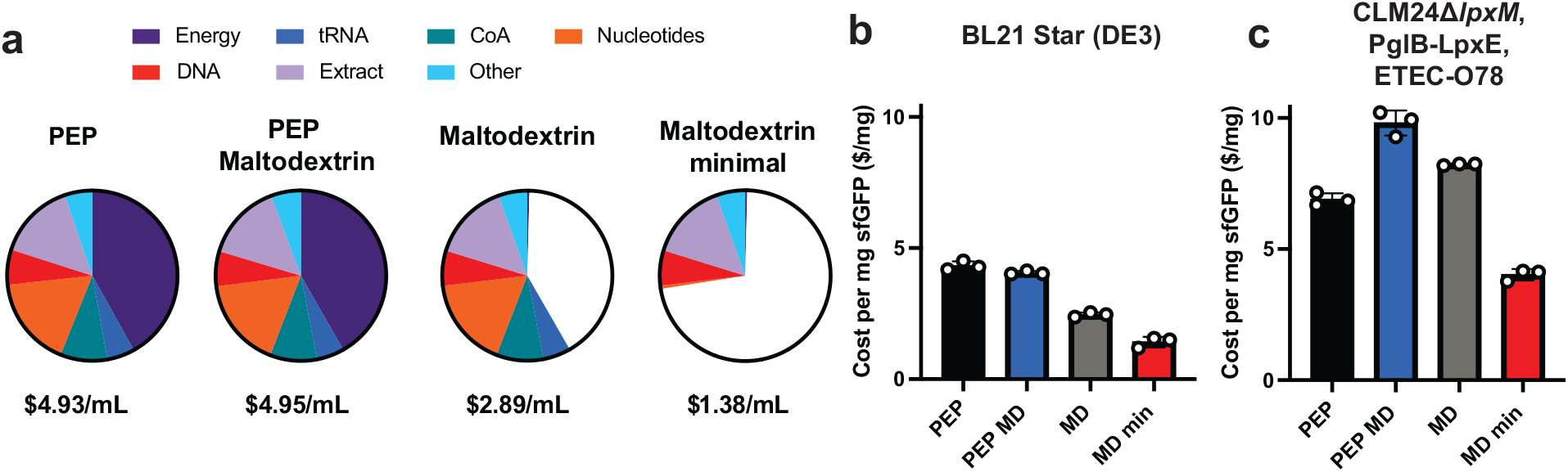
Maltodextrin can be effectively used as both an energy source and lyoprotectant for low-cost CFE. (A) Cost per mL CFE reaction was calculated for each formulation: PEP with no lyoprotectant, PEP with maltodextrin supplemented as a lyoprotectant (PEP MD), maltodextrin as both energy source and lyoprotectant (MD), and maltodextrin without CoA, tRNA, and replacing NTPs with NMPs (MD min). Costs are based upon only raw materials included in the reaction purchased at laboratory scale using calculations in **Supplementary Tables S1-3**. (B) Cost per gram sfGFP in CFE reactions using BL21 Star (DE3) extract in all four formulations. (C) Cost per gram sfGFP in CFE reactions using CLM24 Δ*IpxM* extract in all four formulations. Error bars represent standard deviation of 3 CFE reactions.

To test whether these low-cost maltodextrin formulations could work in practice, we assembled these formulations and evaluated their ability to produce protein. Specifically, we tested four formulations (PEP, PEP+MD, MD, and MD min; **Table S4**) using extracts from BL21 Star (DE3) and a specialized iVAX production strain (CLM24 Δ*IpxM*) harboring glycosylation machinery (**Table S5**)^13^ for the synthesis of sfGFP in fresh reactions. We first optimized the addition of exogenous phosphate in the form of potassium phosphate dibasic (75 mM) and buffer (Bis-Tris and HEPES) in maltodextrin-based reactions. For BL21 Star (DE3) extracts, 57 mM Bis-Tris buffer (pH 10) was optimal and maintained higher final reaction pH (**Figure S2**)^31,44^. Notably, all formulations with these extracts produced roughly the same amount of sfGFP (~1,000 μg/mL), indicating that the removal of reagents did not significantly impact protein yields (**Figure S3A, Figure S4**). Interestingly, the CLM24 Δ*IpxM* performed better with the HEPES buffer at pH 7.2 (**Figure S5**) and 60 mg/mL maltodextrin appeared to have a detrimental impact on sfGFP yields with ~70% protein produced in the PEP MD formulation and ~50% protein produced in both the MD and MD min formulations compared to the original (PEP) formulation (**Figure S3B**). Despite this difference, the MD min formulation has a significantly lower cost per milliliter (**Figure 2B** and **2C**) and enables protein yields sufficient for glycoconjugate vaccine production (~100 μg/mL)^50^, with a maximum yield of ~350 μg/mL sfGFP in the iVAX strain.

### Low-cost CFE formulations retain activity when stored at up to 50 °C

We next sought to evaluate the thermostability of the optimized, low-cost CFE formulations after lyophilization. We lyophilized all four formulations using CLM24 Δ*IpxM* extracts and stored each at room temperature, 37 °C, and 50 °C, for 4 weeks (**Figure 3A).** We rehydrated samples with 5 μL of water and measured maximum initial rates over the first ninety minutes (**Figure 3C, 3E,** and **3G; Figure S6)** as well as endpoint sfGFP concentrations after 20 hours of incubation (**Figure 3B, 3D,** and **3F**). Lyophilization did not reduce activity compared to fresh controls (**Figure S7**), but we found that the supplementation of purified T7 RNA polymerase required for transcription (often stored in glycerol) must be dialyzed to remove glycerol (into S30 buffer, see methods) to maintain activity (**Figure S8**).

**Figure 3.**
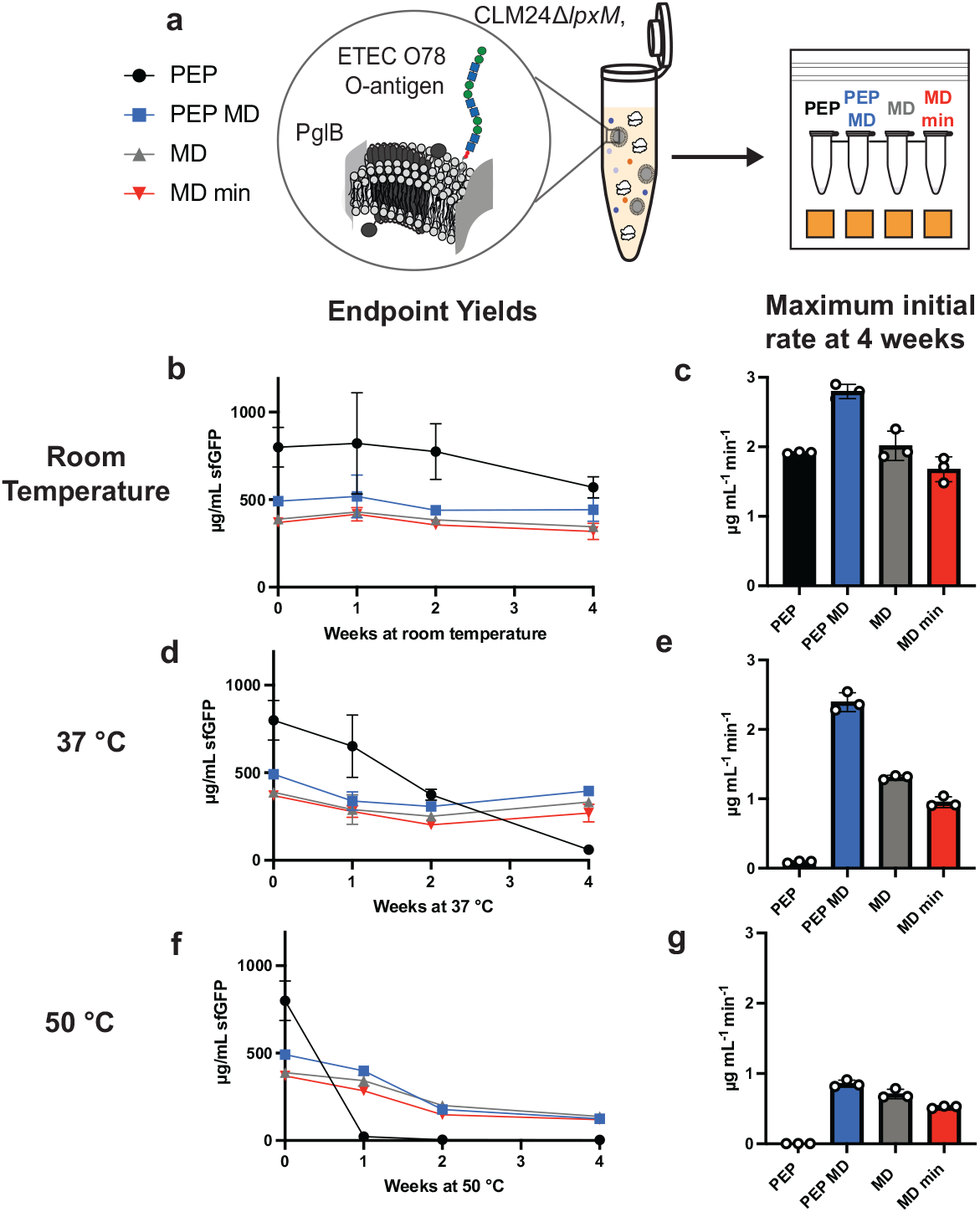
Low-cost formulations preserve CFE reactions with iVAX extract when stored at up to 50 °C. (A) Schematic of CFE reaction storage conditions. After four weeks of storage at room temperature (B-C), 37 °C (D-E), and 50 °C (F-G), lyophilized CFE reactions were rehydrated with 5 μL of water and incubated at 30 °C for 20 hours and endpoint sfGFP yields and maximum initial protein synthesis rates were measured. Error bars represent standard deviation of 3 CFE reactions.

After four weeks of storage at room temperature, all formulations retained activity using CLM24 Δ*IpxM* extracts (**Figure 3B**). However, at elevated temperatures, the PEP-only formulation lost all activity after four weeks of storage at 37 °C (**Figure 3D**) and after one week of storage at 50 °C (**Figure 3F**), while the maltodextrin-containing formulations retained activity, albeit reduced at 50 °C (**Figure 3F**). Interestingly, reactions that use maltodextrin as the energy source have slower initial protein production rates despite similar endpoint yields, suggesting that maltodextrin is more slowly metabolized (**Figure 3C, 3E,** and **3G**). While lyophilized maltodextrin-based formulations have been shown to be stable at ambient conditions^49^, this work demonstrates the first instance, to our knowledge, of high-temperature storage (50 °C) and stability of assembled CFE reactions where the energy substrate is also acting as the lyoprotectant.

### Low-cost, thermostable CFE enables conjugate vaccine production and storage

With a low-cost, thermostable formulation at hand, we wanted to produce and store conjugate vaccines as a potential use case. We have previously shown that coupled CFE and glycosylation (iVAX) reactions are stable at room temperature for up to three months^13^, but higher temperatures are likely encountered during distribution without cold-chain temperature control. To test elevated temperatures on storage of iVAX reactions, we considered a model conjugate vaccine for distribution in resource-limited settings comprised of the O-antigen from enterotoxigenic *E. coli* ETEC O78, a strain of enterotoxigenic *E. coli* responsible for diarrheal disease, conjugated to the licensed carrier protein (PD) from *H. influenzae*^51–53^. We lyophilized iVAX reactions with CLM24 Δ*IpxM* extracts containing the necessary glycosylation machinery and the new CFE formulations. After storage at room temperature (**Figure 4A**), 37 °C (**Figure 4B**), and 50 °C (**Figure 4C**) for one, two, and four weeks, we measured carrier protein (PD) produced via ^14^C-labeled leucine incorporation. Lyophilized reactions behaved similarly under elevated temperatures when producing sfGFP (**Figure 3**) and PD (**Figure 4A-C**).

**Figure 4.**
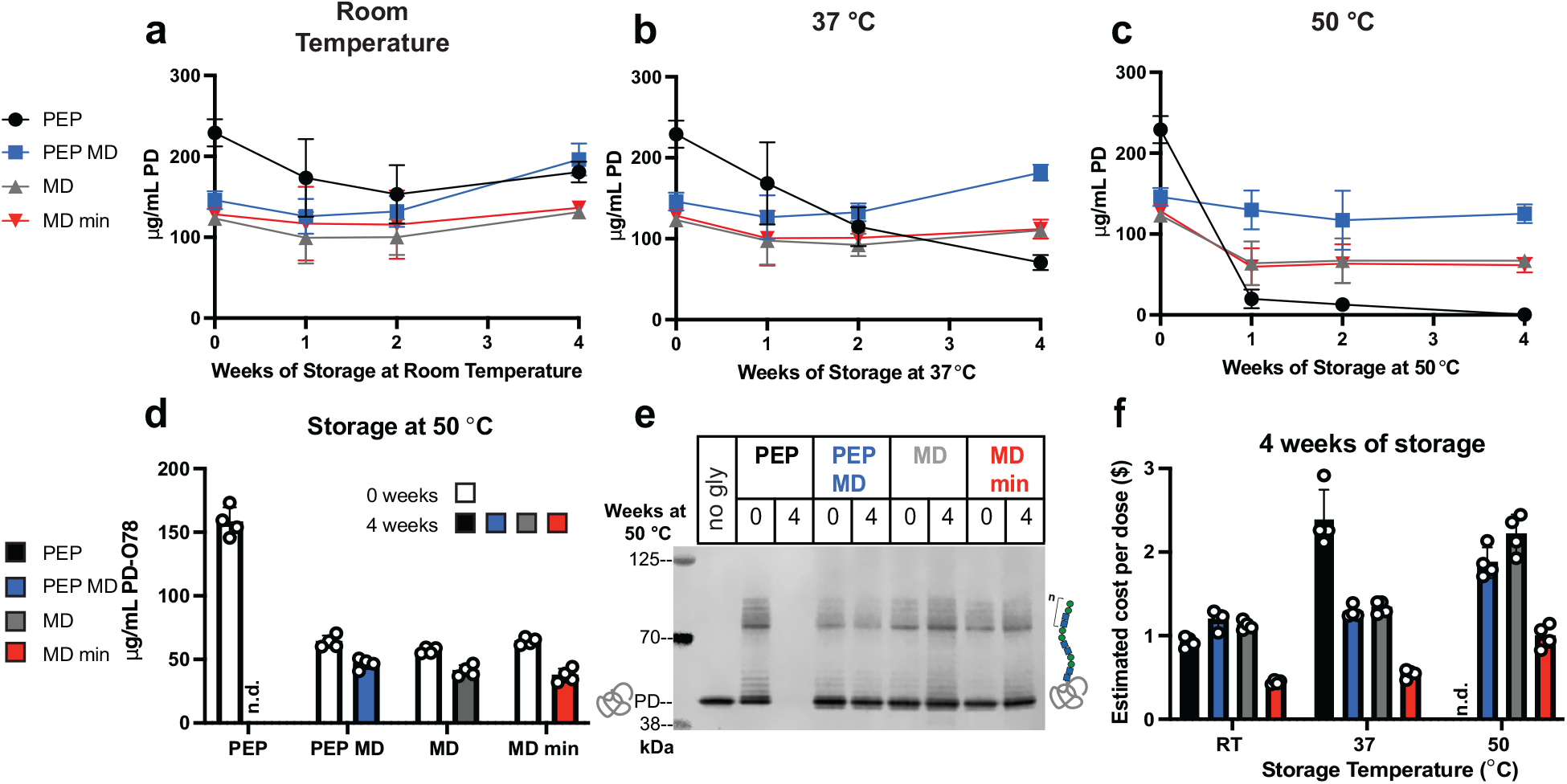
Maltodextrin based formulations with iVAX extract enable production of conjugate vaccine molecules at low cost after high temperature storage. Yields of carrier protein (PD) were measured from lyophilized 15 μL reactions stored for up to 4 weeks at (A) room temperature, (B) 37 °C, and (C) 50 °C. CFE reactions were rehydrated with 15 μL of water and incubated at 30 °C for 20 h. Yields of glycosylated carrier protein (PD) were measured (D) and observed via Western blot (E) for reactions that were stored at 50 °C. (F) Estimated cost per dose of conjugate vaccines produced by CFE reactions stored for 4 weeks at each tested temperature. Error bars represent standard deviation of 4 CFE reactions.

Glycosylation was initiated after 4 hours of protein synthesis (~100-200 μg/mL of carrier protein produced) (**Figure S9A**), yielding > 57 μg/mL of glycosylated PD in all conditions before storage^50^. The glycosylation activity is retained in all formulations before storage (**Figure S10A**) and preserved after four weeks of storage at 50 °C (**Figure 4E, Figure S10C (uncropped)**). Each formulation with protein produced can efficiently glycosylate PD as seen by the characteristic O-antigen banding pattern (varying number of repeated monomers) on the anti-His Western blot (**Figure 4E**). The glycoprotein produced is also cross-reactive with serum specific for the ETEC O78 O-antigen (**Figure S10B**).

While maltodextrin-containing reactions slightly impair glycosylation efficiency, with ~50% glycosylation when MD is present compared to ~70% for PEP formulations (**Figure S10D**), these reactions produce more protein after storage at elevated temperatures, yielding higher concentrations of glycosylated product than the PEP formulations (**Figure 4D**). At 24 μg of conjugate vaccine per dose, we estimated that the MD min formulation could synthesize conjugate vaccines for ~$0.50 per dose after storage at 37 °C for four weeks and ~$1.00 per dose after storage at 50 °C for four weeks (**Figure 4F**), making it the most cost-effective, thermostable cell-free glycoprotein synthesis formulation. Even before storage, the MD min formulation still has a cost benefit due to the significantly cheaper cost of raw materials in the reactions (**Figure S9B**). These developments reduce the cost of iVAX reactions capable of synthesizing conjugate vaccines and enable activity after weeks of storage at elevated temperatures that could be encountered during distribution without cold-chain temperature control^28^.

### Conjugate vaccines produced using the MD min formulation elicit bactericidal antibodies

Finally, we tested the effectiveness of lyophilized PD-O78 conjugates synthesized using the MD min CFE formulation. We scaled up production, purified conjugates, and immunized 8 BALB/c mice with ~24 μg of conjugate or negative control (aglycosylated PD or PBS). Mice were then boosted with 24 μg of conjugate on days 21 and 42, with serum collected on day 56 at the end of the trial (**Figure 5A**). ETEC O78 O-polysaccharide (O-PS) specific antibodies were generated in mice that received purified conjugate derived from lyophilized MD min CFE reactions that was significant over both negative controls tested (**Figure 5B**). We also tested the bactericidal activity of the serum collected from mice and observed ~50% survival for one- and ten-fold serum dilutions for the conjugates that was not observed with sera derived from mice who received the control treatments (**Figure 5C**). Together this data shows that conjugates derived from our new cost-effective, stable, MD min formulation are effective at eliciting bactericidal antibodies against ETEC O78 O-PS. As demonstrated by the robust glycosylation profile observed in our cell-free glycosylation reactions after storage at a variety of temperatures (**Figure 4E**), we expect that conjugate vaccines derived from reactions stored at elevated temperature conditions will remain effective.

**Figure 5.**
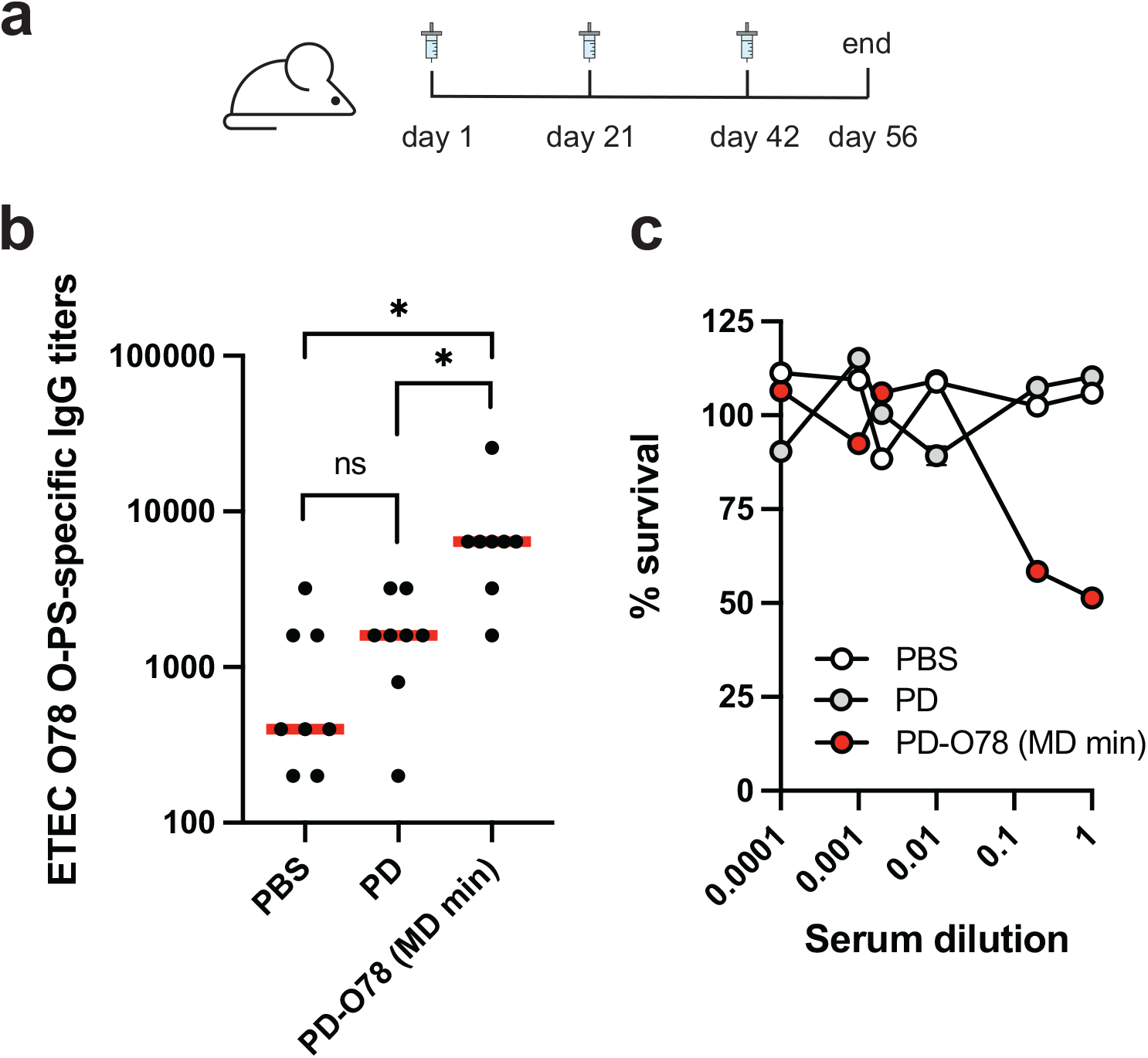
Conjugate vaccines produced using the MD min CFE formulation elicit antibodies that are bactericidal. (A) Lyophilized MD min CFE reactions using iVAX extracts were used to synthesize anti–ETEC O78 conjugate vaccines for immunization studies. Groups of BALB/c mice were immunized subcutaneously with a 1:1 mixture of adjuvant and PBS or ~24 μg of the following cell-free derived immunogens: unconjugated protein D (PD), or PD modified with O78 O-PS from a minimal iVAX reaction (PD-O78 (MD min)). Each group was composed of eight mice. Mice were boosted on days 21 and 42 with identical doses of antigen. (B) ETEC O78 O-PS-specific IgG titers were measured by enzyme-linked immunosorbent assay (ELISA) in endpoint (day 56) serum of individual mice (black dots) with recombinant O-PS immobilized as antigen. Mean titers of each group are also shown (red lines). Statistical significance was calculated by unpaired twotailed *t*-test using GraphPad Prism 9 for MacOS software (version 9.2.0) with a single asterisk (*) indicating *p*-value < 0.05 and ns indicating not significant. (C) Bacterial killing activity of serum antibodies corresponding to the same groups as in (B). Survival data were derived from a standard serum bactericidal assay (SBA) where dilutions of pooled sera from immunized mice were tested against ETEC O78 strain H10407 in the presence of human complement. Values for % survival were determined from the colony forming units (CFUs) counted at each individual serum dilution. Data in C represent average error bars for two independent samples (n = 2).

## Discussion

Cost and stability of CFE reactions are key barriers to point-of-need use, such as iVAX for glycoconjugate vaccine production. Here, we build upon previous cell-free optimizations and studies to identify a low-cost thermostable reaction formulation. A key innovation of this work is the use of maltodextrin to simultaneously stabilize lyophilized reactions at high temperatures and reduce reaction costs. Maltodextrin can be used as energy in the CFE reaction and as a lyoprotectant without extensive optimization. To our knowledge, this is the first characterization of CFE reactions using a non-phosphorylated energy substrate at elevated temperatures (above ambient). We were further able to reduce the cost of the reaction to ~25% of the original, by identifying a maltodextrin minimal (MD min) formulation that is economically beneficial for multiple extract source strains tested and is still capable of synthesizing protein after storage for 4 weeks at 50 °C. This formulation supports protein synthesis in extracts produced from the common high-yielding strain BL21 Star (DE3) as well as a specialized iVAX strain tailored to produce complex glycosylated products.

Successful conjugate vaccine distribution as determined by the MenAfriVac campaign must achieve <$0.50 per dose and tolerate extreme storage temperatures (40°C)^27,28^. Importantly, we show that our low-cost CFE formulations are in line with these metrics and can synthesize effective model conjugate vaccine molecules against ETEC O78 after storage for up to 4 weeks at 37 °C at this price point. Additionally, the formulation is still active after storage at up to 50 °C, although the price increases to $1.00 per dose. In fact, our maltodextrin minimal (MD min) system is capable of synthesizing ~40 μg/mL of glycoconjugate vaccine molecule after storage at all conditions after 4 weeks, higher than previously reported concentrations for this molecule^13^. Importantly, all formulations with maltodextrin retain protein synthesis activity after high temperature storage, while the activity of the original formulation (PEP) declines (37 °C) or disappears (50 °C).

Our formulations achieved <$1.00 per dose for all storage temperatures tested, and the MD min formulation stored at room temperature can reach ~$0.40 per dose. These cost estimates were determined based on raw materials purchased at the laboratory scale^15^. Labor and capital equipment costs required for production are highly dependent on production scale and were therefore not included. We anticipate this work and the continued interest in CFE systems will lead to new metrics to more accurately predict CFE cost at a variety of scales, formal large-scale economic analyses, and further optimization of cell-free reaction formulations to improve commercial feasibility.

Looking forward, this work is an important step in the implementation of cell-free reactions for decentralized manufacturing and builds on past work by taking advantage of the multiple properties of maltodextrin as a reaction component. Importantly, glycosylated products now join other highly sought after molecules that can be produced in lyophilized CFE systems following a range of storage conditions^35^. Taken together, the generation of effective ETEC O78 conjugate vaccines in a low-cost, thermostable formulation advances the iVAX platform and increases accessibility of the technology that can be used to synthesize glycoprotein vaccines in low-resource settings.

## Materials and Methods

### Extract Preparation

Cells were grown in shake flasks at the 1 L scale or in a Sartorius Stedim BIOSTAT Cplus bioreactor at the 10L scale. BL21 Star (DE3) cells were inoculated at optical density at 600 nm (OD_600_) = 0.08 and grown in 2xYTPG at pH 7.2 at 37 °C. Cells were induced at OD_600_ = 0.6 with 0.5 mM of IPTG to induce T7 RNA polymerase expression and harvested at OD_600_ = 3. CLM24 Δ*IpxM* cells transformed with plasmids pSF-PglB-LpxE^13^ and pMW07-O78^13,54,55^ were inoculated at OD_600_ = 0.08 and grown in 2xYTP with no glucose and carbenicillin at 100 μg/mL and chloramphenicol at 34 μg/ml supplemented at 37 °C. Cells were induced at OD_600_ = 0.8-1 with 0.02% arabinose to induce expression of PglB and the ETEC-O78 O-antigen and harvested at OD_600_ = 3. All subsequent steps were performed on ice unless otherwise stated. Cells were harvested by centrifugation at 5,000 x g for 15 minutes and then washed 3 times with S30 buffer (10 mM Tris acetate pH 8.2, 14 mM magnesium acetate, and 60 mM potassium acetate). Following washing, cells were pelleted at 7,000 x g for 10 minutes, then either flash frozen and stored at −80°C, or directly resuspended for lysis.

BL21 Star (DE3) cells were resuspended in 1 mL/g S30 buffer. Cells were then lysed using a Q125 Sonicator (Qsonica, Newtown, CT) with a 3.175 mm diameter probe at a frequency of 20 kHz and 50% amplitude. Energy was delivered to cells in pulses of 10 s followed by 1 s off until 640 J was delivered to each 1-mL aliquot of resuspended cells. Following lysis, cells were centrifuged for 12,000 x g for 10 minutes. Supernatant was then collected, flash frozen and stored at −80 °C as the final extract.

CLM24 Δ*IpxM* cells were resuspended in 1 mL/g S30 buffer. Cells were then homogenized using an EmulsiFlex-B15 (1 L scale) or an EmulsiFlex-C3 (10 L scale) high-pressure homogenizer (Avestin, Inc. Ottawa, ON, Canada) with 1 pass at a pressure of ~21,000 psig. Following lysis, cells were centrifuged for 12,000 x g for 10 minutes. Supernatant was then collected and incubated at 37 °C for 1 hour in a runoff reaction. Cells were then centrifuged once more at 10,000 x g for 10 minutes and then the supernatant was flash frozen and stored at −80 °C as the final extract. Reagents involved in extract preparation are included in **Table S2.**

### Plasmids

All plasmids used in this study are listed in **Table S5**. No new plasmids were cloned in this study, and all appropriate references are cited.

### CFPS Reactions

Reactions were run at the 5-μL scale in PCR tubes in a qPCR instrument set to 30 °C incubation or at the 15-μL scale in 1.5-mL microcentrifuge tubes in a 30 °C incubator (Axygen). Reactions were run for 20 hours when synthesizing sfGFP. Reactions containing lyoprotectants were supplemented with trehalose (Sigma, T0167), sucrose (Sigma, S0389), Dextran 70 (TCI chemicals, D1449), glucose (Sigma, G8270), maltose (Sigma, M9171), or maltodextrin-dextrose equivalent 4.0-7.0 (Sigma,419672), at the appropriate final concentrations (10-100 mg/mL) as described in the text. A fresh stock solution of 300 mg/mL maltodextrin was prepared fresh before reactions were set up and added to CFPS reactions at the appropriate concentration. All other lyoprotectants were prepared and stored at −20 °C.

Reactions for each formulation were prepared as described below and in **Table S4**:

#### PEP

Each reaction was prepared as described previously^50^ unless otherwise noted, to contain 13.33 ng/uL plasmid, 30% (vol./vol.%) S12 extract, and the following: 10 mM magnesium glutamate (Sigma, 49605), 10 mM ammonium glutamate (Biosynth, FG28929), 130 mM potassium glutamate (Sigma, G1501), 1.2 mM adenosine triphosphate (Sigma A2383), 0.85 mM guanosine triphosphate (Sigma, G8877), 0.85 mM uridine triphosphate (Sigma U6625), 0.85 mM cytidine triphosphate (Sigma, C1506), 0.034 mg/mL folinic acid, 0.171 mg/mL *E. coli* tRNA (Roche 10108294001), 2 mM each of 20 amino acids, 30 mM phosphoenolpyruvate (PEP, Roche 10108294001), 0.4 mM nicotinamide adenine dinucleotide (Sigma N8535-15VL), 0.27 mM coenzyme-A (Sigma C3144), 4 mM oxalic acid (Sigma, PO963), 1 mM putrescine (Sigma, P5780), 1.5 mM spermidine (Sigma, S2626), and 57 mM HEPES (Sigma, H3375). T7 was supplemented to reactions at a final concentration of 15-20 μg/mL using the iVAX strain either in 50% glycerol or dialyzed into S30 buffer supplemented with 2 mM DTT.

#### PEP MD

Maltodextrin at a final concentration of 60 mg/mL was supplemented to the PEP reaction formulation described above. See **Table S4** for more details.

#### MD

Maltodextrin at a final concentration of 60 mg/mL was supplemented to the PEP reaction formulation described above and PEP was removed. Potassium phosphate dibasic was supplemented to the PEP reaction formulation at a final concentration of 75 mM unless otherwise noted. Potassium phosphate dibasic (Sigma, 60353) was prepared and pH was adjusted to 7.2 with acetic acid. For BL21 Star (DE3) extract-based reactions, Bis-Tris (Sigma, B9754) with unadjusted pH was added at a concentration of 57 mM and HEPES was removed. See **Table S4** for more details

#### MD min

Reactions were prepared according to the MD reaction formulation described above with the removal of tRNA and CoA. NTPs were also replaced by equal concentration of NMPs (CMP: Sigma C1006, UMP: Sigma U6375, AMP: Sigma 01930, GMP: Sigma G8377). NMPs were prepared at a stock concentration of 0.5 M by dissolving in nuclease free water and pH was adjusted to 7.2 with acetic acid. See **Table S4** for more details.

### Lyophilization and Packaging

CFPS reactions were set up as described above in the CFPS Reactions methods section. Reactions were set up on ice and aliquoted into PCR strip tubes with 1 hole in the lid created by an 18-gauge needle. Samples were kept on ice in aluminum blocks (Cole-Parmer 6361504) and then samples (in blocks) were flash frozen in liquid nitrogen. Frozen samples in blocks were then transferred to a multi-tainer manifold adapter on a VirTis Benchtop Pro Lyophilizer (SP scientific). Lyophilization was performed at 100 mT and a condenser set to −80 °C. Samples were lyophilized overnight for 16-20 hours. Following lyophilization, samples were packaged (all replicates stored together for each tested time and temperature condition) in a FoodSaver bag with 2-4 Dri-Card desiccant cards and then vacuum sealed under ambient conditions with a FoodSaver vacuum sealer. Packaged samples were then stored at room temperature at the bench (~22 °C), or in incubators set to either 37 °C or 50 °C as indicated for the appropriate storage time. Lyophilized controls were rehydrated immediately after removal from the lyophilizer and not stored or packaged in a vacuum sealed bag.

### Protein Quantification

#### For sfGFP measurement

2 μL of CFPS reaction was diluted with 48 μL of nanopure water in a black costar 96 well plate. Fluorescence was read on a plate reader and converted to μg/mL sfGFP using a standard curve with sfGFP measured by C^14^ incorporation.

#### For initial sfGFP synthesis rate measurements

fluorescence was measured every 5 minutes by the qPCR machine. Initial rates were calculated by taking the maximum slope over the first 90 minutes of the cell-free protein synthesis reaction. Using a standard curve, relative fluorescence units measured by the qPCR were converted to μg/mL of sfGFP. To calculate the maximum initial slope over the first 90 minutes, a sliding window of 5 time points was used. For each window, the slope was determined based on a regression line fitting the five time points. This was repeated over the 90 minutes, advancing the starting time point of the window by 1 each time. The maximum initial slope was determined independently for each of the 3 replicates, which were then averaged together to determine the overall average maximum initial slope. This process was completed for each individual reaction condition.

#### For PD synthesis

15-μL reactions containing all reagents except the DNA template were lyophilized and then rehydrated with 15 μL of nuclease free water containing 200 ng of PD-4xDQNAT (or no DNA in control reactions), and 10 μM of C^14^ Leucine (PerkinElmer). Following centrifugation at 16,000 x g for 15 minutes, 5 μL of the soluble fraction of each reaction was treated with 5 μL of 0.5 M KOH for 20 minutes at 37 °C. Following incubation, 4 μL of the sample was added to two filtermats (PerkinLermer Printer Filtermat A 1450-421). After the filtermat dried, one filtermat was washed 3 times for 15 minutes with 5% w/v TCA at 4 °C and once with Ethanol for 10 minutes at room temperature. After the washed filtermat dried, scintillation wax (PerkinElmer MeltiLex A 1450-441) was melted on both mats and counts were measured using a Microbeta2 scintillation counter (PerkinElmer). Background radioactivity was measured in CFGpS reactions with no template DNA and subtracted before calculating protein yields. Fraction of incorporated leucine (washed/unwashed counts) was multiped by the overall leucine concentration in the reaction and the molecular weight of pJL1-PD-4xDQNAT (**Table S5**). The amount of protein produced was determined by dividing this value by the number of leucines present in the protein.

### Cell-Free Glycoprotein Synthesis

For PD synthesis and glycosylation, 15 μL reactions were rehydrated with nuclease free water (Ambion) supplemented with 200 ng of pJl1-PD-4xDQNAT and 10 μM of C^14^ Leucine for a total volume of 15 μL added to the reactions. After rehydration, reactions were incubated for 4 hours at 30 °C. After 4 hours, 0.1 % (wt/vol) DDM and 25 mM MnCl2 were added to each reaction to initiate glycosylation and incubated at 30°C for an additional 16 hours. Before analysis, samples were centrifuged at 16,000 x g for 15 minutes and the soluble fraction was removed. The soluble fraction of each reaction was used to measure yields of the accepter protein PD-4xDQNAT by radioactive counting and to load on western blot to verify glycosylation.

### Western Blotting

Samples were loaded on 4-12% Bis-Tris gels and run with SDS-MOPS running buffer supplemented with NuPAGE antioxidant. Samples were then transferred to Immobilon-P-polyvinylidene difluoride (PVDF) 0.45 μm membranes (Millipore, USA) for 55 minutes at 80 mA per blot using a semi-dry transfer cell. Membranes were blocked for 1 hour at room temperature or overnight at 4 °C in Intercept Blocking Buffer (Licor). Primary antibodies used were anti-His (Abcam, ab1187) at 1:7,500 dilution or anti-ETEC-O78 antigen (Abcam, ab78826) at 1:2,500 dilution in Intercept blocking buffer with 0.2% Tween20 were incubated for 1 hour at room temp or overnight at 4 °C. A fluorescent goat, anti-rabbit antibody GAR-680RD (Licor, 926-68071) was used as the secondary antibody at 1:10,000 dilution in Intercept blocking buffer, 0.2% Tween20 and 0.1% SDS for both anti-His and anti-O78 glycan blots. Blots were washed 6 times for 5 minutes after each blocking, primary, and secondary antibody incubation using 1x PBST. Blots were imaged with Licor Image studio and analyzed by densitometry using Licor Image studio Lite. Fluorescence background was subtracted from each membrane before densitometry was performed.

### Cost Analysis

The cost of each CFPS reaction formulation was estimated using lab scale quantities of reagents from vendors utilized in this study (**Table S3**). Labor and equipment costs are not considered in these estimations. For extract cost estimations, it is assumed that 4 mL of extract are produced per L of cell culture and 30% v/v extract is added to CFE reactions. A “base” extract cost of only the components added to the culture for all strains is considered to make cost estimates more generalizable. The cost of variable components such as inducers and antibiotics are approximately the same for both strains used in this study and are dependent on strain and plasmid used to make extract and are thus neglected (**Table S2**). Glycosylation cofactors are included in the vaccine cost estimates. Vaccine cost estimates assume a 24-μg conjugate vaccine dose and take into account the glycosylation efficiency (amount of PD successfully glycosylated) determined in **Figure S10D. Supplementary Table 1-4** includes references, assumptions and detailed cost calculations for each reaction component.

### Mouse Immunizations

#### Glycoconjugate production

Cell-free glycoprotein synthesis reactions were run as described above using the MD min CFE reaction formulation and were scaled up to 5 mL in 50 mL falcon tubes. Reactions were lyophilized overnight for 16-20 hours and then rehydrated with 5 mL of nuclease free water and incubated at 30 °C for 1 hour. Following 1 hour of protein synthesis, glycosylation was initiated and reactions were incubated at 30°C overnight. The unglycosylated PD negative control was synthesized using the PEP CFE formulation in S30 iVAX extract without the ETEC-O78 pathway overexpressed^13^.

#### Glycoconjugate Purification

CFGpS reactions were centrifuged at 20,000 xg for 10 minutes. The supernatant was then mixed with 0.5 mL of Ni-NTA Agarose resin (Qiagen), equilibrated with 50 mM NaH2PO4, 300 mM NaCl, 10 mM imidazole, per 1 mL of CFE reaction and incubated with agitation for 2-4 hours at 4°C. Purification of His-tagged carrier protein (glycosylated and aglycosylated) was carried out according to manufacturer’s protocol as follows. Following incubation with resin, CFE reaction and resin slurry was loaded onto polypropylene columns (Bio-Rad) and washed 2 times with 6 column volumes of buffer containing 50 mM NaH2PO4, 300 mM NaCl, and 20 mM imidazole. Protein was eluted with 50 mM NaH2PO4, 300 mM NaCl, and 300 mM imidazole. The most concentrated elution fractions were pooled and concentrated to ~2 mg/mL, then dialyzed into sterile endotoxin-free PBS and stored at 4°C. Purification fractions were analyzed on an SDS-PAGE gel and Coomassie stained. Densitometry (Licor ImageStudio) of carrier protein/total protein from SDS-PAGE was used to account for percent purity and multiplied by total A280 protein concentration as measured by nanodrop to determine conjugate concentration for mouse immunizations.

#### Mouse immunizations

Groups of eight 6-week-old female BALB/c mice (Harlan Sprague Dawley) were immunized with 50 μL of sterile PBS (pH 7.4, Fisher Scientific) or formulations containing unconjugated nonacylated protein D (PD) from *Haemophilus influenzae* made using the PEP CFE formulation in S30 iVAX extract without the ETEC-O78 pathway overexpressed, or PD modified with ETEC O78 O-PS made using the MD min CFE formulation (PD-O78 (MD min)). The amount of antigen in each preparation was normalized to ensure that ~24 μg of unmodified protein or conjugate was administered per injection. Purified protein groups formulated in PBS were mixed with an equal volume of Adju-Phos aluminum phosphate adjuvant (InvivoGen) before injection. Each group of mice was immunized subcutaneously with vaccine candidates or controls, then boosted 21 and 42 days after the initial immunization. For antibody titering, blood was obtained on days 0, 35, and 49 via submandibular collection, and at study termination on day 56 via cardiac puncture. For bacterial killing assays, final blood collections for all the mice within each group were pooled. All procedures were carried out in accordance with protocol 2012-0132 approved by the Cornell University Institutional Animal Care and Use Committee.

### Enzyme-linked immunosorbent assay (ELISA)

The plasmid pMW07-O78 encoding the pathway for *E. coli* ETEC O78 O-antigen biosynthesis was used to transform *E. coli* JC8031 competent cells. The resulting cells were used to prepare O78 LPS antigen in house by hot phenol water extraction after DNase I (Sigma) and proteinase K (Invitrogen) treatment, as described elsewhere^56^. Extracted LPS was purified using a PD-10 desalting column packed with Sephadex G-25 resin (Cytiva), and concentration was determined using a purpald assay^57^. 96-well plates (MaxiSorp; Nunc Nalgene) were incubated with 0.5 μg/mL of purified O78 LPS diluted in PBS, pH 7.4, 25 μL/well, at 4 °C overnight. Plates were blocked in blocking buffer overnight at 4 °C with 5% (w/v) nonfat dry milk (Carnation) in PBS, then washed three times with 200 μL PBS-T (PBS, 0.05% Tween 20) per well. Serum samples isolated from the collected blood draws of immunized mice were appropriately serially diluted in triplicate in blocking buffer and added to the plates for 2 hours at 37 °C. Plates were washed three times with PBS-T (+ 0.03% BSA (w/v)), then incubated for 1 hour at 37 °C in the presence of a horseradish peroxidase–conjugated antibody, goat anti-mouse IgG (Abcam, 1:25,000 dilution). After three PBS-T + 0.3% BSA washes, 50 μL of 3,3’-5,5’-tetramethylbenzidine substrate (1-Step Ultra TMB-ELISA; Thermo Fisher Scientific) was added to each well, and the plates were incubated at room temperature in the dark for 30 min. The reaction was stopped by adding 50 μL of 2 M H2SO4, and absorbance was measured at a wavelength of 450 nm using a FilterMax F5 microplate spectrophotometer (Agilent). Serum antibody titers were determined by measuring the lowest dilution that resulted in signals that were 3 standard deviations above the background controls of no serum. Statistical significance was determined in GraphPad Prism 9 for MacOS (Version 9.2.0) using an unpaired two-tailed *t*-test.

### Serum bactericidal assay (SBA)

A modified version of a previously described SBA method was followed^54^. ETEC H10407 cells were grown overnight from a frozen glycerol stock, then seeded 1:20 in Luria Bertani (LB) medium. Log-phase grown bacteria were harvested, adjusted to an OD_600_ of 0.1, then further diluted 1:5,000 in Hanks’ Balanced Salt Solution with 0.5% bovine serum albumin (BSA) (Sigma Aldrich). Assay mixtures were prepared in 96-well microtiter plates by combining 20 μL of serially diluted heat-inactivated test serum (with dilutions ranging from 1-10^4^), and 10 μL of diluted bacterial suspension. After incubation with shaking for 60 minutes at 37°C, 10 μL of active or inactive complement source was added to each well, to a final volume percent of 25%. Heat-inactivated complement was prepared by thawing an aliquot of active pooled human complement serum (Innovative Research, ICSER1ML), incubating in a 56°C water bath for 30 minutes, and cooling at room temperature. Assay plates were incubated with shaking at 37 °C for 60–90 minutes, then 10 μL was plated from each well (diluted to 50 μL in LB) on LB agar plates. Serum samples were tested and plated in duplicate, and colonies were counted (Promega Colony Counter) after 16–18 hours of incubation at 30 °C. Colony forming units (CFUs) were counted for each individual serum dilution, and SBA titers were determined by calculating percent survival at various serum dilutions. Data were plotted as percentage survival versus serum dilution.

## Acknowledgments

We acknowledge Jessica Stark and Jasmine Hershewe for helpful discussions and contribution of cell-free reagents used in this work and Kosuke Seki for the development of the code used for maximum initial rate calculations. This work was supported by the Bill and Melinda Gates Foundation (OPP1217652 to M.P.D. and M.C.J.), Defense Threat Reduction Agency (HDTRA1-15-10052 and HDTRA1-20-10004 to M.P.D. and M.C.J.), Army Contracting Command (W52P1J-21-9-3023), and National Science Foundation (CBET-1936823 to M.P.D. and CBET – 1936789 to M.P.D. and M.C.J.). K.F.W acknowledges the National Defense Science and Engineering (NDSEG) Fellowship Program (ND-CEN-013-096) sponsored by the Army Research Office. M.C.J. thanks the David and Lucile Packard Foundation. A.W. acknowledges support from the Cornell Postdoctoral Scholars Program. D.A.W. acknowledges support from the National Science Foundation Graduate Research Fellowship under grant no. DGE-1842165.

## Author Contributions

All of the authors designed the research; K.F.W., A.W., D.A.W, S.E.S., P.D., and J.L. performed research; K.F.W., A.W., D.A.W, and S.E.S analyzed data; Y.F.-C., M.P.D. and M.C.J. directed research; and K.F.W, A.S.K., and M.C.J. wrote the paper. All authors reviewed and edited the paper.

## Competing Interests

M.P.D. has a financial interest in Gauntlet, Inc., Glycobia, Inc., SwiftScale Biologics, Inc., Versatope, Inc., Gauntlet Bio, and UbiquiTx, Inc. M.C.J. has a financial interest in SwiftScale Biologics, Gauntlet Bio, Pearl Bio, Inc., Design Pharmaceutics, and Stemloop Inc. M.P.D.’s and M.C.J.’s interests are reviewed and managed by Cornell University and Northwestern University, respectively, in accordance with their competing interest policies. All other authors declare no competing interests.

