## Supplementary Information for "A low-cost, thermostable, cell-free protein synthesis platform for on demand production of conjugate vaccines"

#### Supplementary Figures 1-10
#### Supplementary Tables 1-5
#### Supplementary References

### Supplementary Figures

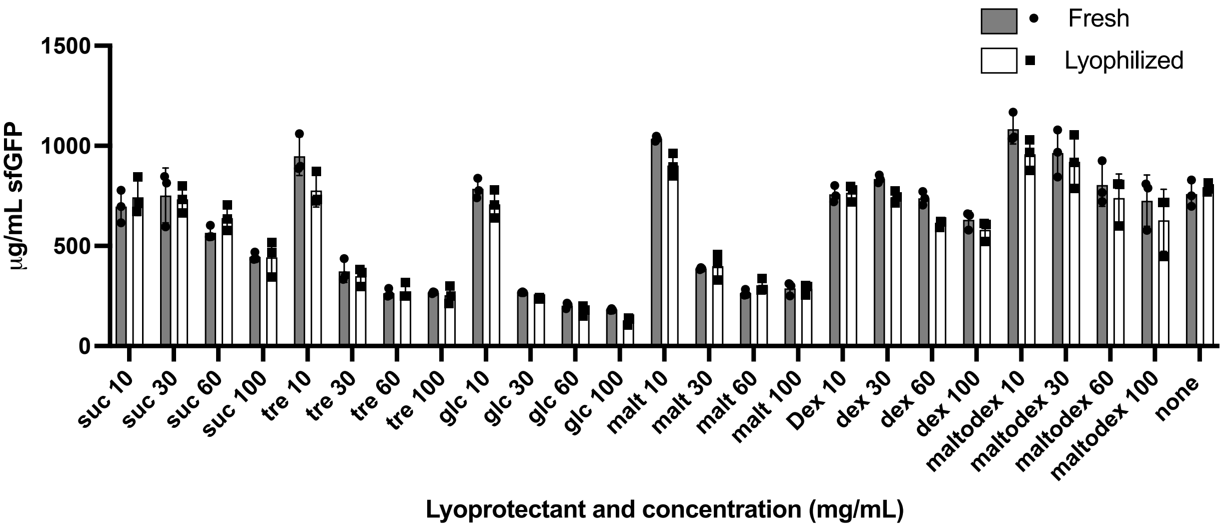

**Figure S1. Impact of lyoprotectant additives on fresh and lyophilized CFE reactions with BL21 Star (DE3) extract.** The impact of sucrose (suc), trehalose (tre), glucose (glc), maltose (malt), dextran (dex), and maltodextrin (maltodex) at 0, 10, 30, 60, 100 mg/mL final concentration on fresh (grey bars) and rehydrated lyophilized (white bars) CFE reaction productivity. Control with no lyoprotectant (none) is shown on the right. Error bars represent standard deviation of three CFE reaction replicates (n=3).

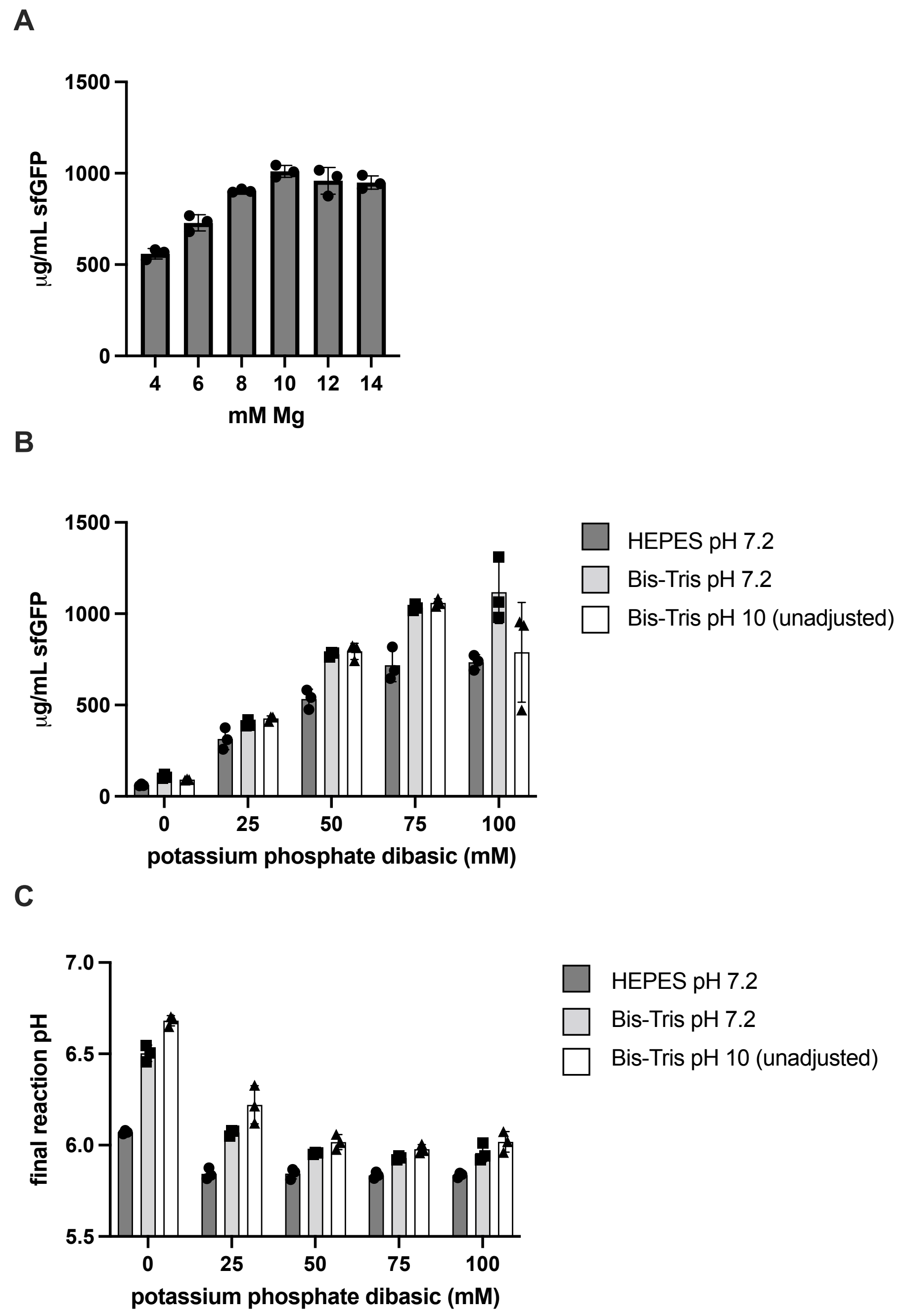

**Figure S2. Optimization of CFE reagents for MD formulation in BL21 Star (DE3) extract.** (A) Magnesium optimization of lysate using PEP as an energy source with sfGFP synthesis as a reporter. (B) Using 10 mM $Mg^{2+}$ in the MD reaction formulation, sfGFP production was used to determine the optimal concentration of potassium phosphate dibasic (pH 7.2) in the CFE reaction. Impact of buffer on the MD formulation was also tested using 57 mM of either HEPES with pH adjusted to 7.2 (dark grey), Bis-Tris with pH adjusted to 7.2 (light grey), or Bis-Tris with unadjusted pH (pH 10) (white). (C) Impact of conditions in B on final pH of the cell-free protein expression measured after 20 hours of sfGFP synthesis at 30 °C. Error bars represent standard deviation of three CFE reaction replicates (n=3).

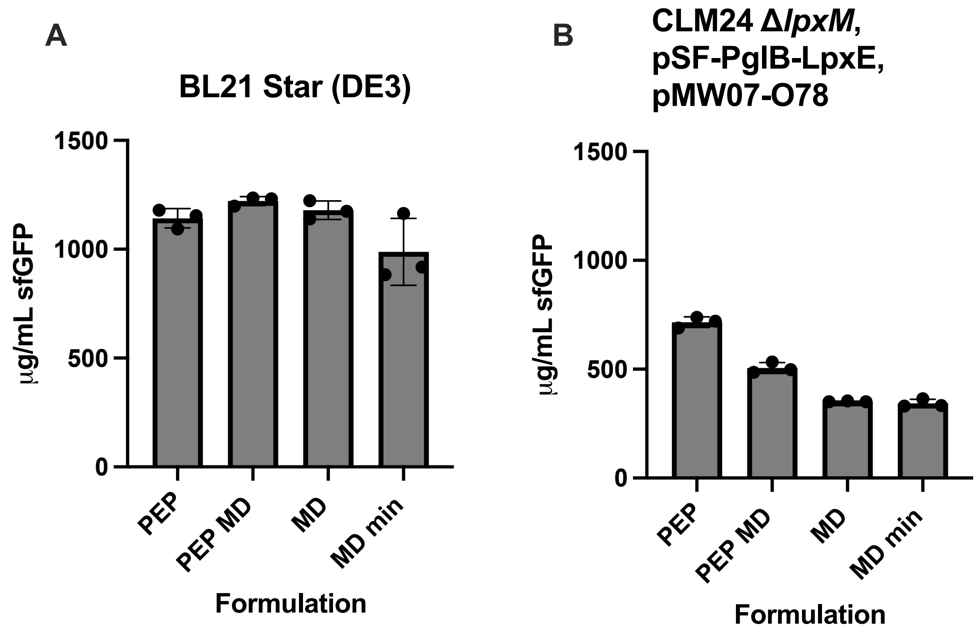

**Figure S3. sfGFP yields of each formulation in CFE reactions using different extracts.** (A) sfGFP synthesis in CFE reactions using BL21 Star (DE3) extract for each formulation. (B) sfGFP synthesis in CFE reactions using the iVAX strain (CLM24 Δ*lpxM* with overexpression of glycosylation machinery from pSF-PglB-LpxE and pMW07-O78 plasmids) extract for each formulation. Error bars represent standard deviation of three CFE reaction replicates (n=3).

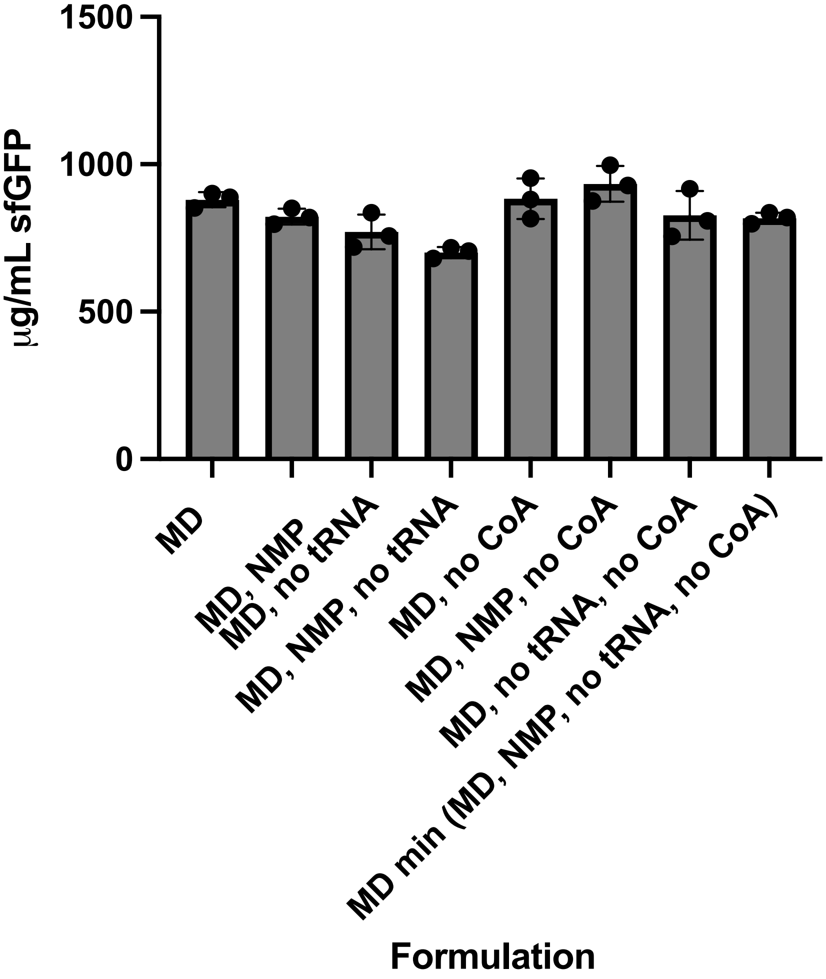

**Figure S4. sfGFP yields in CFE reactions using BL21 Star (DE3) extract showing the impact of the formulation changes made to the MD formulation to arrive at the MD min formulation.** sfGFP yields in a BL21 Star (DE3) extract for the MD formulation (far left) and the impact of changing NTPs to NMPs, removing tRNA, and removing CoA, as well as every combination of the other reagents. sfGFP yields in the MD min formulation (all changes at the same time) are displayed on the far right. Error bars represent standard deviation of three CFE reaction replicates (n=3).

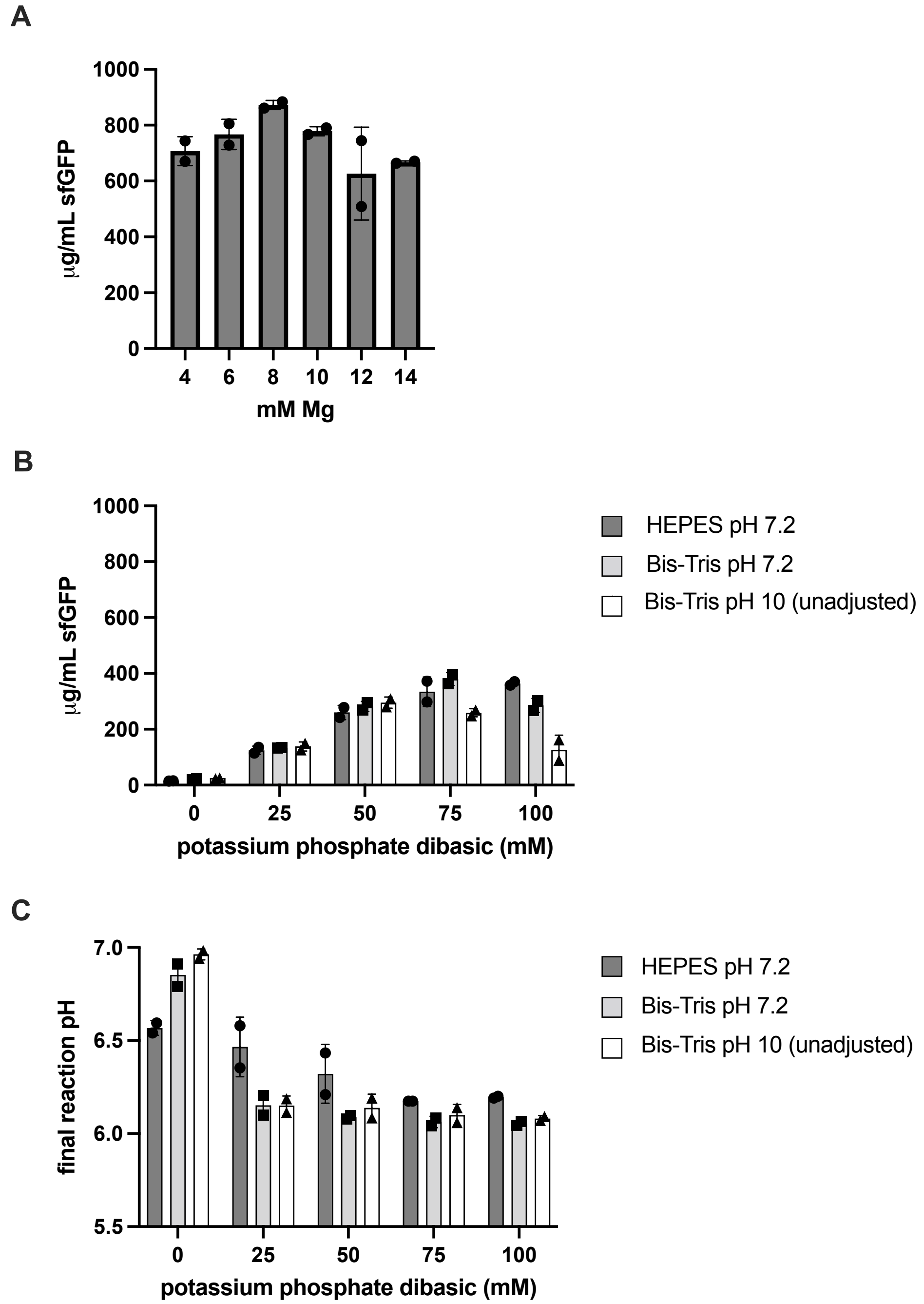

**Figure S5. Optimization of CFE reagents for MD formulation in iVAX extract (CLM24 Δ*lpxM* with overexpression of glycosylation machinery from pSF-PglB-LpxE and pMW07-O78 plasmids).** (A) Magnesium optimization of lysate using PEP as an energy source and sfGFP synthesis as a readout. (B) Using 8 mM $Mg^{2+}$ in the MD reaction formulation, sfGFP production was used to determine the optimal concentration of potassium phosphate dibasic (pH 7.2) in the CFE reaction. Impact of buffer on the MD formulation was also tested using 57 mM of either HEPES with pH adjusted to 7.2 (dark grey), Bis-Tris with pH adjusted to 7.2 (light grey), or Bis-Tris with unadjusted pH (pH 10) (white). (C) Impact of conditions in B on final pH of the cell-free reaction measured after 20 hours of sfGFP synthesis at 30 °C. Error bars represent standard deviation of two CFE reaction replicates (n=2).

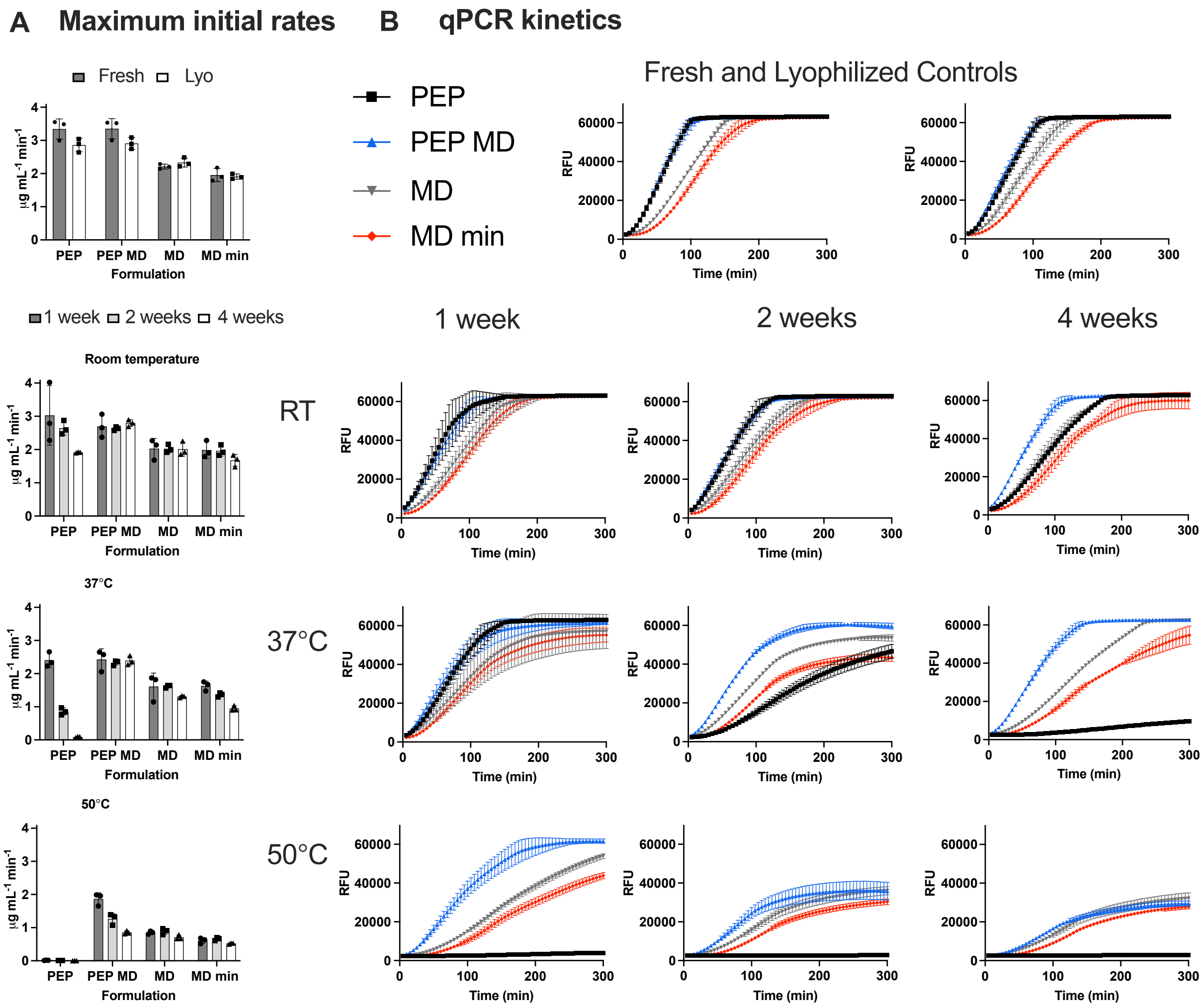

**Figure S6. Rates of sfGFP synthesis in all formulations using the iVAX extract.** (A) Maximum initial rates calculated over the first 90 minutes of protein synthesis using qPCR measurement of fluorescence every 5 minutes for fresh, lyophilized, and 1, 2, and 4 weeks at room temperature, 37 °C, or 50 °C storage. Fluorescence was converted from RFU to µg/mL sfGFP using standard curves with $^{14}$C-labeled sfGFP. (B) qPCR traces recording fluorescence every 5 minutes for all reaction conditions measured in RFUs over the initial 300 minutes of the reaction. While reactions were incubated in the instrument at 30 °C for 20 hours to read endpoint sfGFP, the instrument reaches the limit of detection at ~60,000 RFU, thus only initial kinetic data is shown, and maximum reached at ~60,0000 RFU is due to limit of detection of the instrument and is not representative of protein synthesis levels at that timepoint. Error bars represent standard deviation of three CFE reaction replicates (n=3).

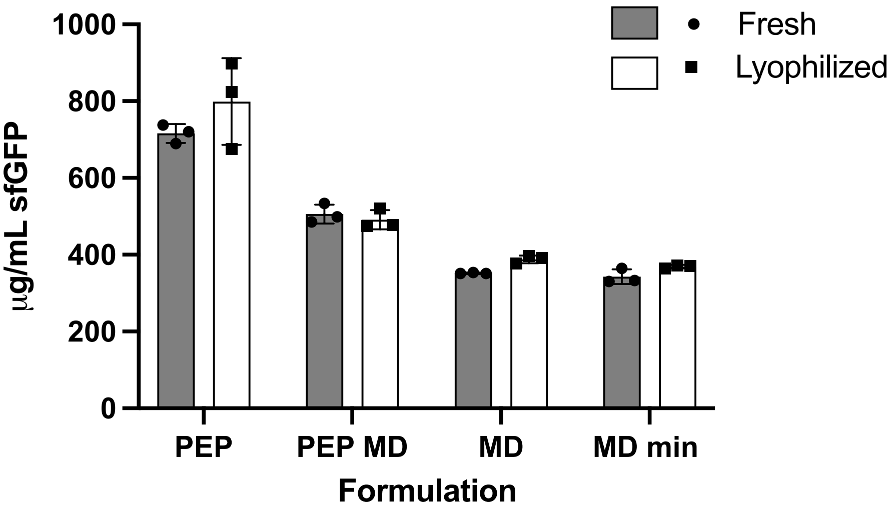

**Figure S7. sfGFP yields of fresh and lyophilized (un-stored/zero-week timepoint) controls of all CFE reaction formulations with the iVAX extract.** sfGFP yields of 5 µL fresh reactions that were not lyophilized (grey) and rehydrated, lyophilized reactions that were un-stored (white) after 20 hours of incubation at 30 °C using the iVAX extract. Error bars represent standard deviation of three CFE reaction replicates (n=3).

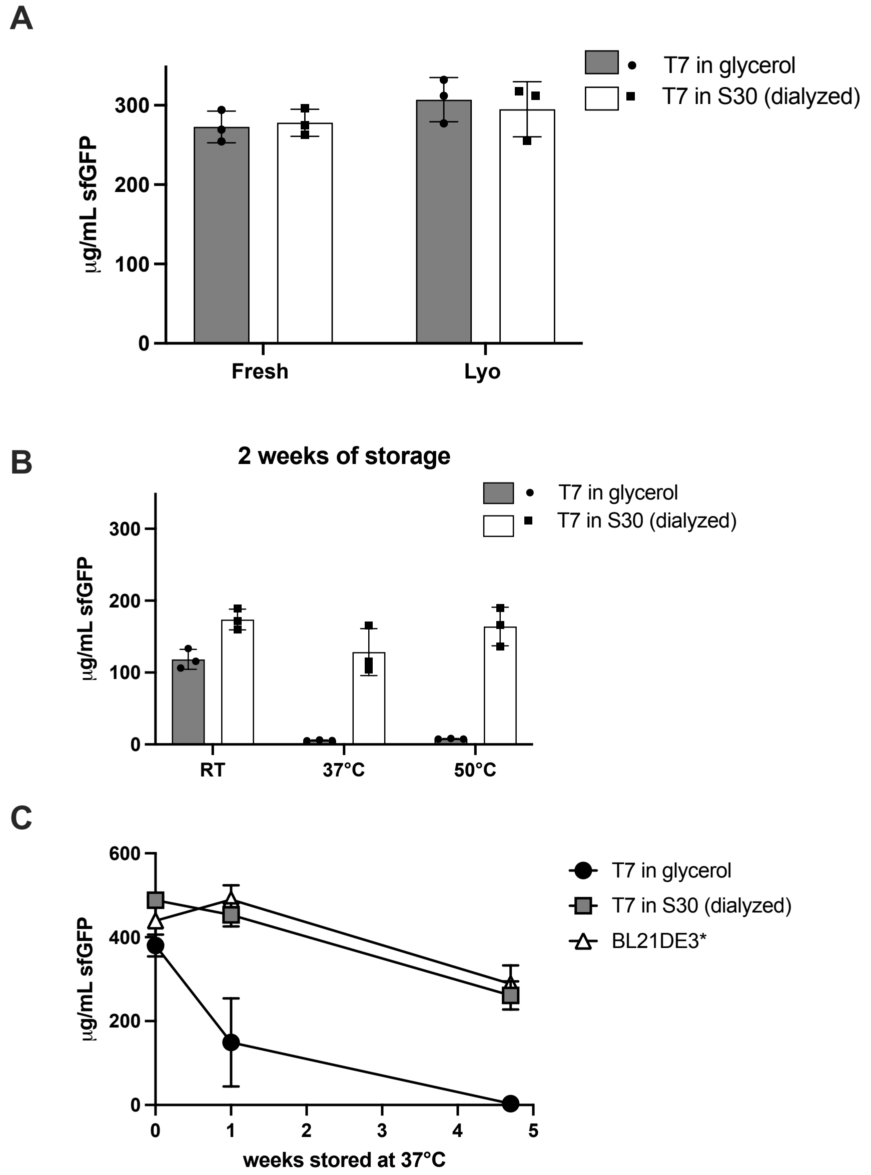

**Figure S8. Impact of glycerol (contained in purified T7) on MD formulation.** (A) sfGFP yields of 5 µL fresh reactions and rehydrated, unstored, lyophilized reactions after 20 hours of incubation at 30 °C using the MD formulation in the iVAX extract. Grey bars have T7 source in 50% glycerol and white bars have the same original T7 source dialyzed into S30 buffer. (B) Reactions from the same experiment shown in A after storage at either room temperature (RT), 37 °C, or 50 °C for 2 weeks. Reactions containing glycerol from the T7 stock are shown in grey while reactions with no glycerol using the dialyzed T7 stock are shown in white. (C) Endpoint sfGFP synthesis timecourse from MD formulation CFE reactions with T7 source as either T7 in glycerol (black circle), T7 in S30 buffer, dialyzed from glycerol (grey squares), or supplementing the reaction with a final concentration of 3.3% v/v BL21 Star (DE3) extract that had T7 overexpressed in the strain before lysis (white triangles). Lysate for this experiment was derived from the parental CLM24 strain not modified for glycosylation or remodeled endotoxin. Reaction conditions optimized for this strain were 8 mM $Mg^{2+}$, 50 mM phosphate, and Bis-Tris pH 10. Error bars represent standard deviation of three CFE reaction replicates for A, B, and C (n=3).

A

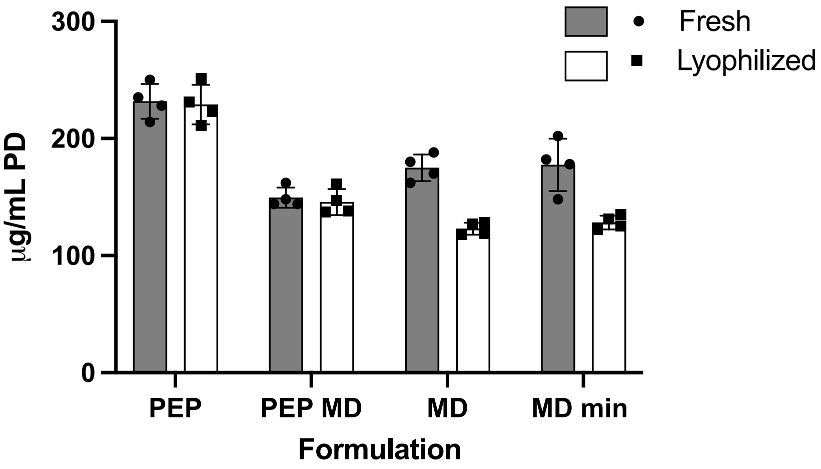

B

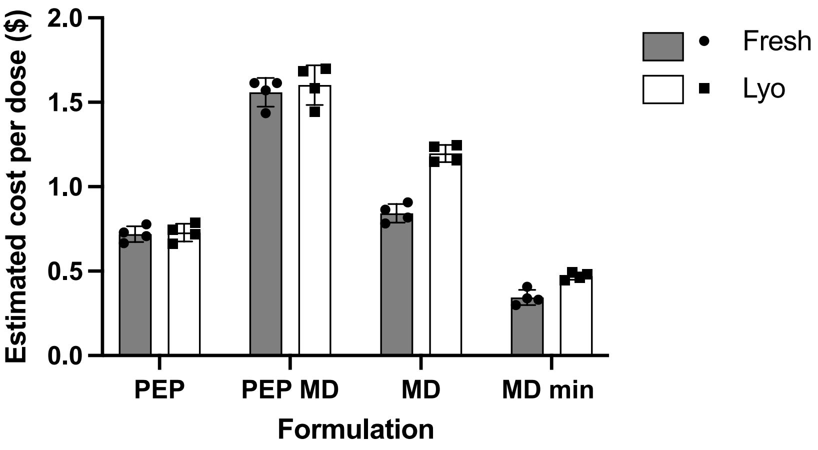

**Figure S9. PD yields and estimated cost per conjugate vaccine dose of fresh and lyophilized (un-stored) controls of all CFE reaction formulations with the iVAX extract.** (A) PD yields of 15 µL fresh reactions that were not lyophilized (grey) and rehydrated, lyophilized reactions that were un-stored (white) after 20 hours of incubation at 30 °C using the iVAX extract. Yields were measured using $^{14}$C-leucine incorporation. (B) Estimated cost per dose of conjugate vaccine obtained from fresh (grey) and lyophilized (white) iVAX reactions. Calculations consider estimated % glycosylation as measured by densitometry in Figure S10D and assume a 24 µg dose. Error bars represent standard deviation of four CFE reaction replicates (n=4).

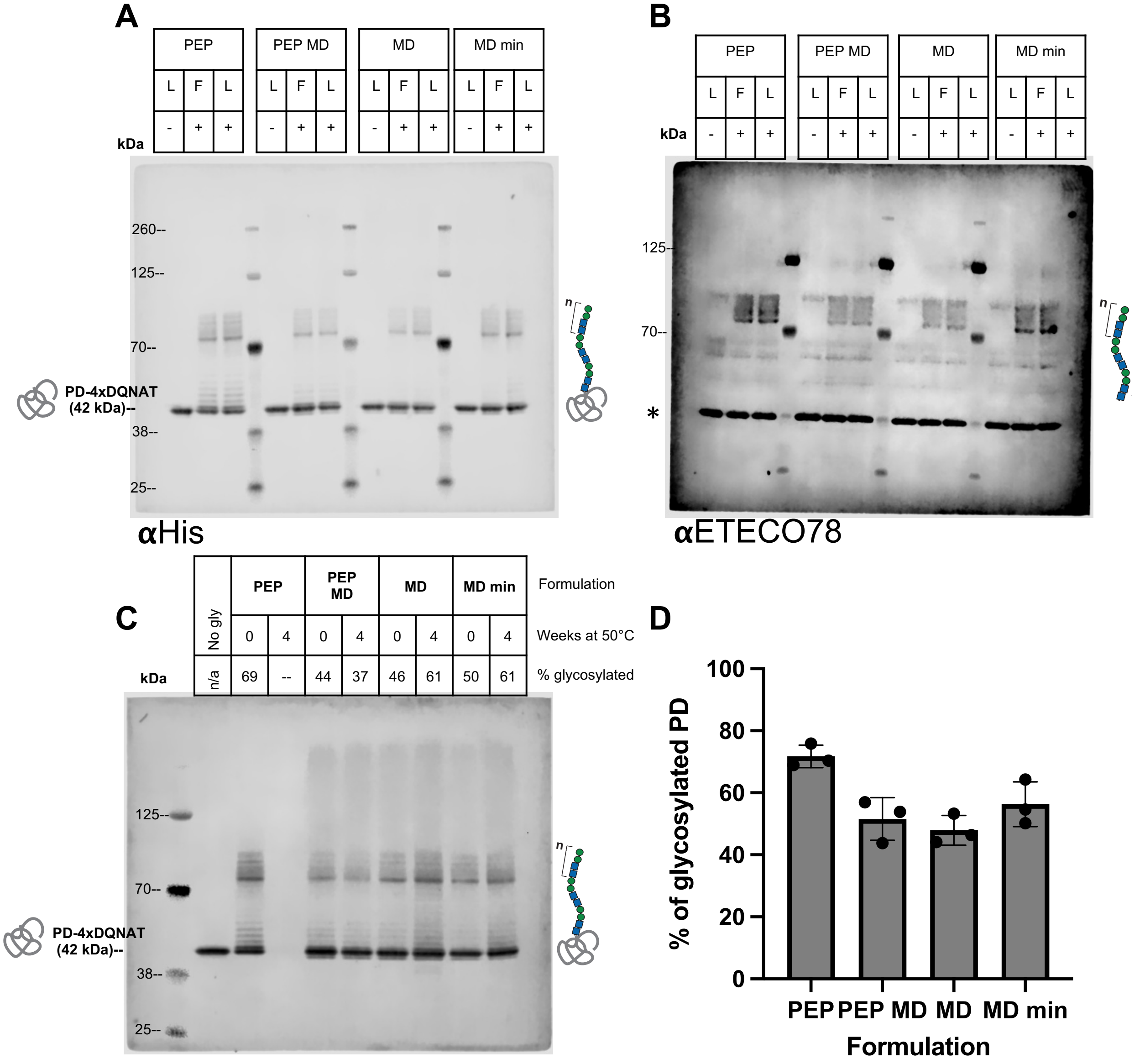

**Figure S10. Glycosylation of PD with ETEC-O78 O-antigen in iVAX reactions.** (A) Anti-His Western blot against His-tagged carrier protein (PD) demonstrating glycosylation with the ETEC O78 O-antigen in fresh and lyophilized reactions. From left to right, three control reactions are shown for each formulation (PEP, PEP MD, MD, and MD min). First a negative lyophilized control (aglycosylated PD) using an iVAX lysate with no ETEC-O78 expression is denoted as (L, -). Then, a fresh, unlyophilized control in the iVAX lysate (glycosylated PD) is denoted as (F, +). Finally, a lyophilized control in the iVAX lysate (glycosylated PD) is shown and denoted as (L, +). Equal concentration of PD as determined by $^{14}$C-leucine incorporation was loaded in each well. Each formulation is separated by a Chameleon 800 ladder annotated on the left-hand side of the blot. (B) Anti-ETEC-O78 glycan western blot demonstrating glycosylation of PD with ETEC-O78 O-antigen in fresh and lyophilized controls. Sample orientation is the same as in A. O-antigen banding is visible between the 70-kDa and 125-kDa MW markers. The band annotated with an asterisk on the left of the blot is a contaminating band found in both glycosylated and unglycosylated samples also observed in our previous work[1]. Equal concentration of PD as determined by $^{14}C$-leucine incorporation was loaded in each well. Each formulation is separated by a Chameleon 800 ladder annotated on the left-hand side of the blot. (C) Uncropped Anti-His western blot shown in Figure 4E demonstrating glycosylation of PD with ETEC-O78 O-antigen in samples stored for 0 weeks (lyophilized, unstored) and samples stored for 4 weeks at 50 °C. The first lane has a negative glycosylation control run with an iVAX lysate with no ETEC-O78 expression in the PEP formulation that was lyophilized and then rehydrated. Then from left to right for each formulation PEP, PEP MD, MD, and MD min, there is a lyophilized sample stored for 0 weeks (lyophilized, unstored) and then a sample that was stored for 4 weeks at 50 °C. Equal concentration of PD as determined by $^{14}C$-leucine incorporation was loaded in each well. For the PEP formulation after 4 weeks storage at 50 °C, no protein synthesis was detected, so the volume equivalent to that used for the least concentrated sample was run on the gel as a verification. Percent glycosylation as estimated by densitometry with Image Studio Lite (Licor) software for each lane is reported above the blot and used for glycoprotein calculation in main Figure 4D. (D) Percent of glycosylated PD (glycosylated PD/total PD) was estimated for each formulation using densitometry and Image Studio Lite (Licor) software. Error bars represent standard deviation of three CFE reactions (n=3). Triplicate values include % glycosylation from fresh and lyophilized samples from A and lyophilized samples from C for each formulation. Gels are representative of three independent experiments. Values were used for cost calculation in main Figure 4F.

### Supplementary Tables

**Supplementary Table 1. Cost breakdown of the CFE reaction formulations used in this work.** Cost per liter of CFE reaction for all reagents present in the PEP, PEP MD, MD, and MD min formulation. This table is based on BL21 Star (DE3) extract as is presented in Figure 2 of the main text.

| Component | PEP ($/L) | PEP MD ($/L) | MD ($/L) | MD min ($/L) |
|---|---|---|---|---|
| magnesium glutamate | 0.94 | 0.94 | 0.94 | 0.94 |
| ammonium glutamate | 5.97 | 5.97 | 5.97 | 5.97 |
| potassium glutamate | 8.48 | 8.48 | 8.48 | 8.48 |
| ATP | 15.16 | 15.16 | 15.16 | |
| CTP | 241.05 | 241.05 | 241.05 | |
| UTP | 277.74 | 277.74 | 277.74 | |
| GTP | 320.63 | 320.63 | 320.63 | |
| AMP | | | | 6.91 |
| CMP | | | | 8.86 |
| UMP | | | | 5.85 |
| GMP | | | | 3.70 |
| folinic acid | 22.62 | 22.62 | 22.62 | 22.62 |
| tRNA | 255.00 | 255.00 | 255.00 | |
| amino acids | 100.54 | 100.54 | 100.54 | 100.54 |
| PEP | 2065.42 | 2065.42 | | |
| maltodextrin | | 19.80 | 19.80 | 19.80 |
| NAD | 110.39 | 110.39 | 110.39 | 110.39 |
| CoA | 435.19 | 435.19 | 435.19 | |
| oxalic acid | 0.19 | 0.19 | 0.19 | 0.19 |
| putrescine | 0.74 | 0.74 | 0.74 | 0.74 |
| spermidine | 5.35 | 5.35 | 5.35 | 5.35 |
| HEPES | 6.06 | 6.06 | | |
| Bis-Tris | | | 9.77 | 9.77 |
| potassium phosphate dibasic | | | 3.18 | 3.18 |
| plasmid DNA | 333.25 | 333.25 | 333.25 | 333.25 |
| extract | 728.25 | 728.25 | 728.25 | 728.25 |
| **Total $/L CFE reaction** | **4932.97** | **4952.77** | **2894.24** | **1374.79** |
| **Total $/mL CFE reaction** | **4.93** | **4.95** | **2.89** | **1.37** |

**Supplementary Table 2. Cost breakdown of cost of cell extract.** Note that the base extract cost is used in all calculations in this work to make claims more generalizable, as variable components are approximately the same for both strains used in this study and are dependent on strain and plasmid used to make extract. Assumptions used in calculations are listed below.

**Assumptions:**

1. 4 mL of extract are produced per liter cell culture.
2. 30% v/v extract is used in CFE reactions.
3. Only raw materials added to cell culture are considered for a base case of extract without variable components such as inducers or antibiotics.
4. Labor costs associated with extract production are not considered.
5. Equipment costs are not considered.

| **Compound Name** | **Vendor** | **Catalog Number** | **$/g** | **$/L culture** |
|---|---|---|---|---|
| **Constant components** | | | | |
| tryptone | Sigma | T7293-1kg | 0.272 | 4.352 |
| yeast extract | Sigma | Y1625-1kg | 0.246 | 2.46 |
| sodium chloride | Sigma | S3014-5kg | 0.0438 | 0.219 |
| potassium phosphate, monobasic | Sigma | P9791-1kg | 0.168 | 0.504 |
| potassium phosphate, dibasic | Sigma | 60353-1kg | 0.312 | 2.184 |
| **Variable components** | | | | |
| glucose | Sigma | G8270-5kg | 0.024 | 0.432 |
| IPTG (0.5 mM) | Sigma | I6758-10g | 49.2 | 5.86218 |
| arabinose (0.02 wt% in media) | Sigma | A3256-1kg | 1.77 | 0.354 |
| carbenicillin disodium salt | Sigma | C1389-10g | 70.7 | 7.07 |
| chloramphenicol | Sigma | C0378-100g | 1.66 | 0.05644 |

| **Extract** | **$/L cells** | **$/mL extract** | **$/mL CFE reaction** | **$/L CFE reaction** |
|---|---|---|---|---|
| **Base** | 9.71 | 2.4275 | 0.72825 | 728.25 |
| **BL21 Star (DE3)** | 16 | 4 | 1.2 | 1200 |
| **iVAX** | 17.19 | 4.2975 | 1.28925 | 1289.25 |

**Supplementary Table 3. Information on all reagents added to the CFE reaction.** Costs are recorded as of January 2022 at lab scale from vendors used in this work.

| Compound Name | Vendor | Catalog Number | $/g | $/L reaction |
|---|---|---|---|---|
| **For PEP formulation** | | | | |
| magnesium glutamate (10 mM) | Sigma | 49605-250g | 0.242 | 0.94 |
| ammonium glutamate | Biosynth | FG28929-.1kg | 3.6383 | 5.97 |
| potassium glutamate | Sigma | G1501-1kg | 0.321 | 8.48 |
| ATP | Sigma | A2383-25G | 22.92 | 15.16 |
| CTP | Sigma | C1506-1g | 538 | 241.05 |
| UTP | Sigma | U6625-1g | 594 | 277.74 |
| GTP | Sigma | G8877-1g | 721 | 320.63 |
| folinic acid | Sigma | 47612-1g | 754 | 22.62 |
| tRNA | Sigma | 10109550001-.5g | 1500 | 255.00 |
| amino acids (cost for 1 g of each) | Sigma | LAA21-1kt | 457 | 100.54 |
| PEP | Sigma | 10108294001-1g | 334 | 2065.42 |
| NAD | Sigma | N8535-15VL | 416 | 110.39 |
| CoA | Sigma | C3144-1g | 2100 | 435.19 |
| oxalic acid | Sigma | P0963-500g | 0.26 | 0.19 |
| putrescine | Sigma | P5780-25g | 4.6 | 0.74 |
| spermidine | Sigma | S2626-25g | 24.56 | 5.35 |
| HEPES | Sigma | H3375-5kg | 0.446 | 6.06 |
| Plasmid DNA | Zymo | D4201-50 preps | 25000 | 333.25 |
| **For modified formulations** | | | | |
| AMP | Sigma | 01930-25g | 14.72 | 6.91 |
| CMP | Sigma | C1006-5g | 28.4 | 8.86 |
| UMP | Sigma | U6375-10g | 18.7 | 5.85 |
| GMP | Sigma | G8377-100g | 10.7 | 3.70 |
| maltodextrin | Sigma | 419672-500g | 0.33 | 19.80 |
| Bis-Tris | Sigma | B9754-1kg | 0.819 | 9.77 |
| potassium phosphate, dibasic | Sigma | 60353-1kg | 0.312 | 3.18 |
| **For iVAX reactions** | | | | |
| DDM | Anatrace | D310S-25g | 39.56 | 39.56 |
| $MnCl_2$ | Sigma | 221279-500G | 0.272 | 1.35 |

**Supplementary Table 4. Components of the CFE reaction formulations used in this work.** Final concentration present of each reagent used in the CFE reaction for all formulations is provided in mM unless otherwise noted in the table. Cells filled in grey indicate that a component is not present in the reaction formulation described by that column. *Choice of buffer is extract strain dependent for the MD and MD min formulations, but both buffers were used at the same final concentration.

| Component | PEP | PEP MD | MD | MD min |
|---|---|---|---|---|
| magnesium glutamate | 10 | 10 | 10 | 10 |
| ammonium glutamate | 10 | 10 | 10 | 10 |
| potassium glutamate | 130 | 130 | 130 | 130 |
| ATP | 1.2 | 1.2 | 1.2 | |
| CTP | 0.85 | 0.85 | 0.85 | |
| UTP | 0.85 | 0.85 | 0.85 | |
| GTP | 0.85 | 0.85 | 0.85 | |
| AMP | | | | 1.2 |
| CMP | | | | 0.85 |
| UMP | | | | 0.85 |
| GMP | | | | 0.85 |
| folinic acid | 0.03 mg/mL | 0.03 mg/mL | 0.03 mg/mL | |
| tRNA | 0.17 mg/mL | 0.17 mg/mL | 0.17 mg/mL | |
| amino acids | 2 | 2 | 2 | 2 |
| PEP | 30 | 30 | | |
| maltodextrin | | 60 mg/mL | 60 mg/mL | 60 mg/mL |
| NAD | 0.4 | 0.4 | 0.4 | 0.4 |
| CoA | 0.27 | 0.27 | 0.27 | |
| oxalic acid | 4 | 4 | 4 | 4 |
| putrescine | 1 | 1 | 1 | 1 |
| spermidine | 1.5 | 1.5 | 1.5 | 1.5 |
| HEPES/Bis-Tris* | 57 | 57 | 57 | 57 |
| potassium phosphate dibasic | | | 75 | 75 |
| plasmid DNA | 13.33 ng/µL | 13.33 ng/µL | 13.33 ng/µL | 13.33 ng/µL |
| extract | 30 % v/v | 30 % v/v | 30 % v/v | 30 % v/v |

**Supplementary Table 5. Strains and plasmids used in this study.**

| Strain or Plasmid | Description |
|---|---|
| BL21 Star (DE3) | *E. coli* B strain for expression. |
| CLM24 Δ*lpxM*[1] | *E. coli* K-12 strain CLM24 with a knockout of the aceyltransferase *lpxM* to alter endotoxin structure. |
| pJL1-sfGFP[2] | sfGFP variant with a C-terminal strep tag in the pJL1 expression vector, |
| pJL1-PD-4x DQNAT[1] | *H. influenzae* protein D modified with a C-terminal 4x DQNAT glycosylation sequon and a 6x His tag, recognized by *C. jejuni* PglB in the pJL1 expression vector. |
| pMW07-O78[1,3,4] | *E. coli* O78 O-antigen gene cluster in the pMW07 expression vector. |
| pSF-PglB-LpxE[1] | *C. jejuni* PglB with a C terminal LpxE phosphatase from *F. tularensis* and a 1x-FLAG tag in the pSF expression vector. |
